# SpinalTRAQ: A novel volumetric cervical spinal cord atlas identifies the corticospinal tract synaptic projectome in healthy and post-stroke mice

**DOI:** 10.1101/2024.08.23.609434

**Authors:** Katherine Poinsatte, Matthew Kenwood, Dene Betz, Ariana Nawaby, Apoorva D Ajay, Wei Xu, Erik J Plautz, Xiangmei Kong, Denise M O Ramirez, Mark P Goldberg

## Abstract

Descending corticospinal tract (CST) connections to the neurons of the cervical spinal cord are vital for performance of forelimb-specific fine motor skills. In rodents, CST axons are almost entirely crossed at the level of the medullary decussation. While specific contralateral axon projections have been well-characterized using anatomic and molecular approaches, the field currently lacks a cohesive imaging modality allowing rapid quantitative assessment of the entire, bilateral cervical cord projectome at the level of individual laminae and cervical levels. This is potentially important as the CST is known to undergo marked structural remodeling in development, injury, and disease. We developed SpinalTRAQ (<u>S</u>pinal cord <u>T</u>omographic <u>R</u>egistration and <u>A</u>utomated <u>Q</u>uantification), a novel volumetric cervical spinal cord atlas and machine learning-driven microscopy acquisition and analysis pipeline that uses serial two-photon tomography-images to generate unbiased, region-specific quantification of the fluorescent pixels of anterograde AAV-labeled CST pre-synaptic terminals. In adult mice, the CST synaptic projectome densely innervates the contralateral hemicord, particularly in laminae 5 and 7, with sparse, monosynaptic input to motoneurons in lamina 9. Motor pools supplying axial musculature in the upper cervical cord are bilaterally innervated. The remainder of the ipsilateral cord has sparse labeling in a distinct distribution compared to the contralateral side. Following a focal stroke of the motor cortex, there is a complete loss of descending corticospinal axons from the injured side. Consistent with prior reports of axon collateralization, the CST spinal projectome increases at four weeks post-stroke and continues to elevate by six weeks post stroke. At six weeks post-stroke, we observed striking synapse formation in the denervated hemicord from the uninjured CST in a homotopic distribution. Additionally, CST synaptic reinnervation increases in the denervated lamina 9 in nearly all motoneuron pools, exhibiting novel patterns of connectivity. Detailed level- and lamina-specific quantification of the bilateral cervical spinal cord synaptic projectome reveals previously undescribed patterns of CST connectivity in health and injury-related plasticity.

## Introduction

Corticospinal (CST) neurons control dexterous fine motor tasks, such as reaching and grasping, in rodents, non-human primates, and humans (Alstermark & Isa, 2012; Heffner & Masterton, 1975; Lemon, 2008). The mammalian CST that originates in the primary motor (M1) and somatosensory cortices passes through the internal capsule and crosses at the pyramidal decussation to innervate the contralateral spinal cord throughout the cervical spinal cord (C1-C8) (Watson, 2021). Primate and rodent CST anatomy differ in several key aspects, most notably in the trajectory of the descending tract (dorsal column in rodents versus lateral and ventral columns in primates and humans) (Lacroix et al., 2004; Lemon, 2019; Nudo & Masterton, 1990). In rodents, CST almost entirely crosses at the medullary pyramids with collaterals predominantly innervate contralateral interneurons in laminae 4-8, (Kaiser et al., 2019; Ueno et al., 2012) and, unlike primates, are reported to provide little-to-no direct input to motoneurons in healthy adult mice (Alstermark & Ogawa, 2004; Gu et al., 2017; Lemon, 2019; Lemon, 2008). The CST targets premotor neuronal populations with distinct biological functions in discrete anatomical regions (Azim et al., 2014; Bui et al., 2013; Hayashi et al., 2018; Levine et al., 2014; Ni et al., 2014; Ronzano et al., 2022; Ueno et al., 2018).

The CST undergoes extensive remodeling in development (Canty & Murphy, 2008; Martin et al., 2007; Welniarz et al., 2017) and after central nervous system injury (Chen et al., 2022; Kaiser et al., 2019; Liu et al., 2021; Ueno et al., 2012) and disease (Lemon, 2021; Zdunczyk et al., 2018). Notably, during development, as opposed to adulthood, the CST differentially targets lower motor neurons directly via an auxiliary branch in the ventrolateral funiculus that is quickly pruned (Gu et al., 2017, Gu et al., 2019, Gu et al., 2020). Injury in the primary motor cortex radically alters the CST, inducing degeneration of the contralateral CST and denervation of the cervical spinal cord (Chen et al., 2022; Kaiser et al., 2019). In response to this unilateral denervation, the healthy CST tract, originating from uninjured motor cortex, undergoes dynamic remodeling within the spinal cord (Carmichael et al., 2017; Chen et al., 2022; Liu et al., 2013). Our lab and others have demonstrated a significant increase in axon length and density of midline-crossing CST fibers (Chen et al., 2022; Kaiser et al., 2019; Lapash Daniels et al., 2009; Liu et al., 2009; Ueno et al., 2012). This axonal remodeling in the denervated cord is behaviorally consequential (Liu et al., 2009), as the number of midline-crossing fibers correlates with the recovery of fine motor skills, and silencing newly crossing CST fibers reinstates motor deficits in recovered animals after stroke (Bachmann et al., 2014; Wahl et al., 2014; Wiessner et al., 2003).

While CST axon pathways are well characterized in certain rodent and primate models, this information does not fully describe the final connectivity pattern, since CST axons can pass through gray matter regions without interacting synaptically. A projectome based on regional quantification of presynaptic terminals adds important detail regarding final projection targets as well as the relative density of synaptic contacts. We developed SpinalTRAQ (<u>S</u>pinal cord <u>T</u>omographic <u>R</u>egistration and <u>A</u>utomated <u>Q</u>uantification), a novel pipeline for 3-D fluorescence image acquisition and analysis of the CST synaptic projectome of the mouse cervical spinal cord. Building on our established whole brain image analysis and registration pipeline (Ortega et al., 2020; Poinsatte et al., 2019; Ramirez D. M. O., 2019), SpinalTRAQ utilizes fluorescent viral tract-tracing, serial two-photon tomography (STPT), and custom cluster computing workflows to create an end-to-end pipeline for 3D visualization, atlas registration, machine learning-driven image segmentation and automated quantification of cervical spinal cord volumetric datasets. In this study, we assess spatial dynamics of the bilateral cervical CST projectome and demonstrate marked regional and temporal remodeling in the weeks after focal stroke of the unilateral motor cortex. These findings introduce a new, adaptable pipeline for spinal cord imaging and analysis, and highlight its utility in elucidating synapse-level changes in CST projections in health and disease.

## Results

### SpinalTRAQ: Spinal cord Tomographic Registration and Automatic Quantification pipeline for region-specific quantification of the cervical spinal cord

The SpinalTRAQ pipeline consists of (1) volumetric serial two-photon tomography (STPT) imaging of cervical spinal cords with endogenous fluorescent signals of interest, (2) parallel post-processing including (a) registration of STPT images into a custom cervical spinal cord atlas and (b) supervised machine learning-driven pixel classification of endogenous fluorescent signals of interest, and (3) automated region-specific quantification of classified pixels (Fig. 1). To establish the pipeline, we traced the CST synaptic projectome in adult male C57/BL6 mice by injecting 1 uL of SynaptoTag4 into the M1 cortical area, which induced expression of eGFP in CST pre-synaptic terminals and tdTomato in descending CST axons (Fig. 2A, B) (Li et al., 2021). Volumetric whole cervical spinal cord datasets were generated using STPT imaging, including imaging at 3 optical depths, and images were subjected to the analysis pipeline containing three main components: pixel classification, atlas registration, and quantification of classified pixels in 3D anatomical space. cervical cord image volumes were registered to a novel mouse cervical spinal cord reference atlas (Fig. 2B, C) (Fiederling et al., 2021a, 2021b). Using this newly established pipeline, we generated a probability map of classified pixels in registered 3-D atlas space for spatially sensitive quantification of the CST synaptic projectome in healthy, adult mice (Supplemental Video 4, 5 and 6). Using SpinalTRAQ, we quantified the probability of classification of a pixel as a pre-synaptic terminal within each of 94 volumetric regions defined at each of the eight cervical levels (Fig. 2D).

**Figure 1:**
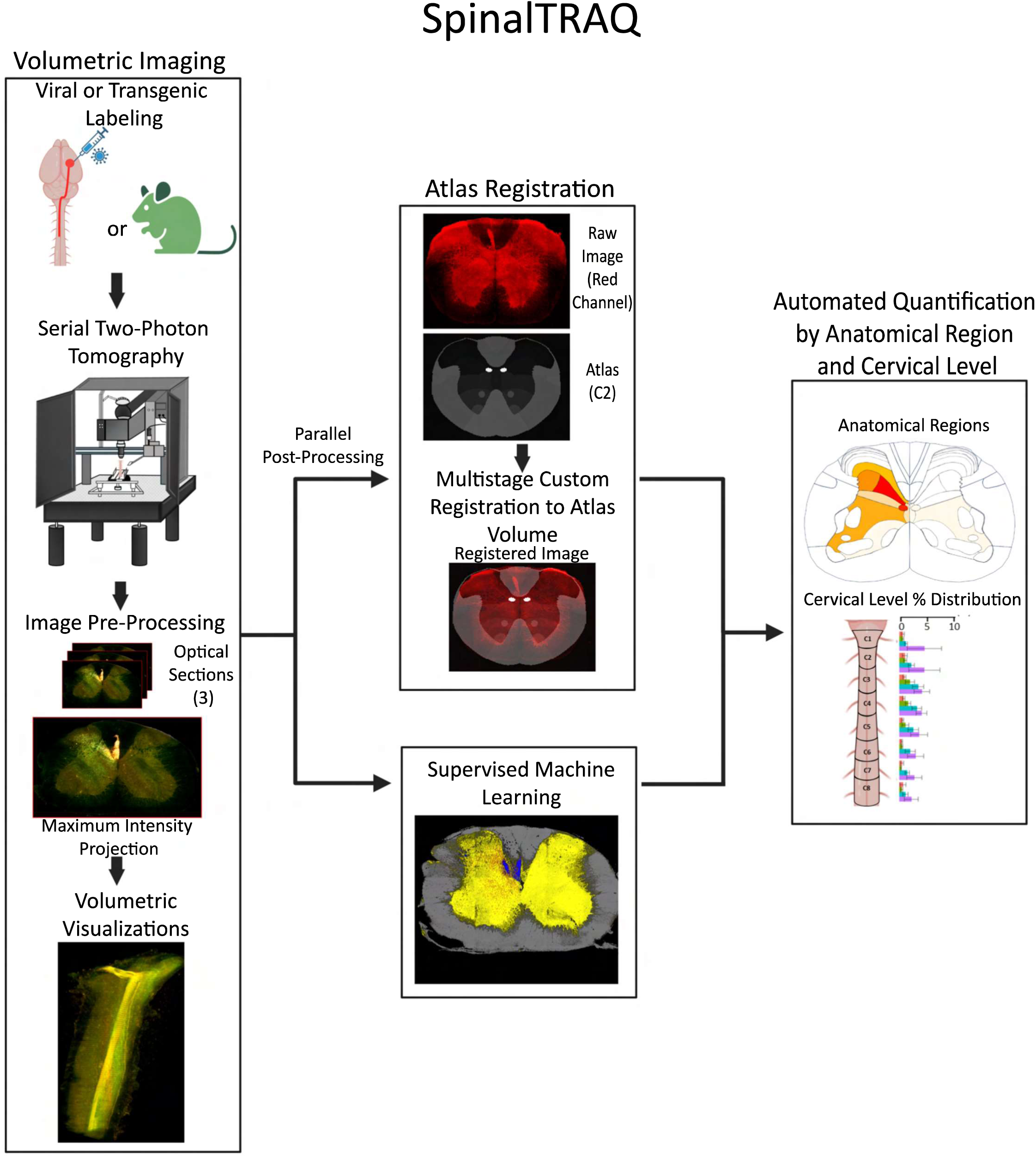
SpinalTRAQ: Application of a digitally annotated spinal cord atlas and custom automated pipeline for unbiased region-specific quantification in volumetric image stacks of mouse cervical spinal cord acquired via STPT. Graphical depiction of SpinalTRAQ workflow. Left, Mouse spinal cord is fluorescently labeled using viral or transgenic methods and imaged using STPT (TissueCyte 1000). Images are pre-processed, and visualized in 3D. Middle, parallel computational workflows allow automated registration of cervical cord volumes into a digital atlas and pixel classification of relevant fluorescent signals via supervised machine learning. Right, Registration of pixel classified images (i.e. probability maps) into the reference atlas and custom scripts are used to generate quantitative results on the location of pre-synaptic terminals or other fluorescent signals of interest across cervical cord laminae and levels.

**Figure 2:**
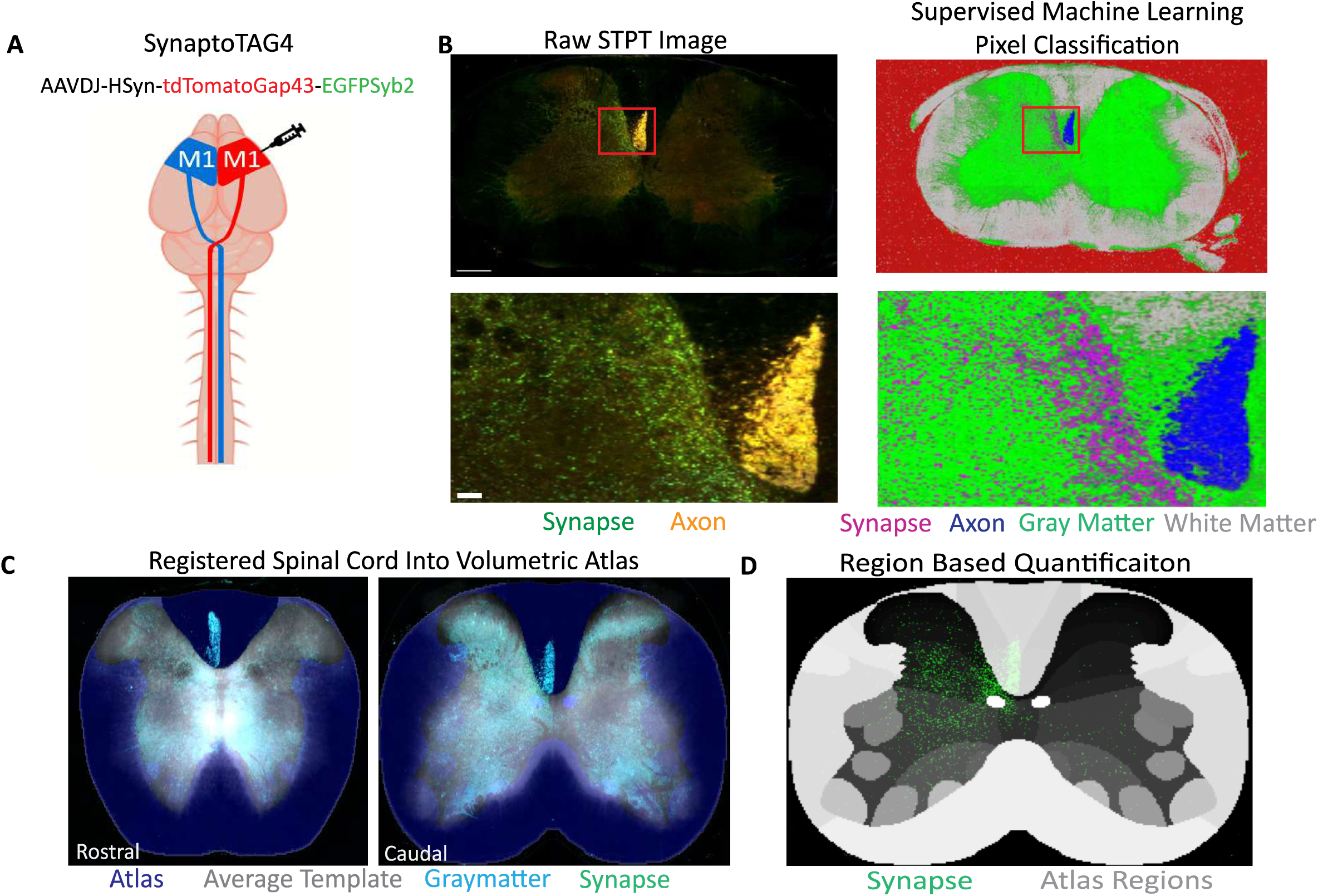
Classifying the CST synaptic projectome in the whole mouse cervical spinal cord. (A) Graphical representation of experimental design, showing a unilateral motor cortex injection of the AAV SynaptoTag4 (AAVDJ-HSyn-tdTomatoGap43-EGFPSyb2-WPRE) into healthy adult mice. (B) Maximum intensity projection of 3 full resolution raw optical planes of the entire cord section (upper left panel, 400um scale bar) with zoomed in section of laminae 4/5 (lower left panel showing red boxed area, 50um scale bar). GFP+ pre-synaptic terminals are shown in green and dual GFP and tdTomato positive axons residing in the CST are shown in yellow. Right panels show the output of supervised machine learning classification via ilastik identifying pre-synaptic terminals/synapses (magenta), axons (blue), gray matter (green), white matter (gray), and background (red) in the same image. (C) Spinal cord autofluorescence (cyan) registered into the average template (grey) at C3(left) and C7 (right). (D) Representative example of a single image’s positive GFP pixel probability map (green) in digital space with the region annotation overlay (gray).

We assessed the accuracy of SpinalTRAQ region classifications using transgenic mice expressing GFP under the choline acetyltransferase (ChAT) promoter (Supplemental Figure 1A). ChAT expression is limited to a prominent large population of spinal motoneurons in lamina 9 and some sparse interneuron populations (Gotts et al., 2016; Shneider et al., 2009). 92.11% of labeled pixels were classified within lamina 9 (n=3) (Supplemental Fig. 1B, Supplemental Fig. 1D). A small proportion of classified soma GFP+ pixels were localized to laminae 7 and 8 neighboring regions to lamina 9 (Supplemental Fig. 1B).

### The level- and lamina-specific CST spinal synaptic projectome in healthy, adult mice

We characterized the CST pre-synaptic terminal distribution (i.e., the CST synaptic projectome) in uninjured mice to provide a comprehensive quantitative analysis of the bilateral CST in whole cervical spinal cords. Mice received a unilateral injection of Synaptotag4 in the right motor cortex 2 weeks prior to sacrifice. TdTomato+ axons were visualized in previously reported locations including the contralateral dorsal column, contralateral dorsolateral funiculus, and more sparsely, and highly variable between animals, in the ipsilateral dorsal column and ipsilateral ventrolateral funiculus (data not shown).

SpinalTRAQ assessment of eGFP+ presynaptic terminals revealed dense innervation within the contralateral cervical gray matter (93.2 ± 2.25% of total labeled synapses, n=4 mice) (Fig. 3A-C). In agreement with axon tracing experiments, eGFP+ terminals were much more sparsely distributed in the ipsilateral cervical hemicord (2.57 ± 3.80%, n=4 mice). (Fig. 3C). CST projections distributed differently at different cervical levels, ranging in the contralateral cord from 7.5% (level C1) to 16.1% (level C5) (Fig. 3C). Synaptic terminals in the ipsilateral hemicord were also similarly distributed, ranging from 0.43 to 0.96% (Fig. 3C). Interestingly, while CST synaptic projections predominantly targeted cervical levels C5-C7 in the contralateral hemicord (42.88 ± 2.65%), cervical level C2 contained the greatest percentage of CST pre-synaptic terminals in the ipsilateral hemicord (0.96 ± 0.425% ipsilateral vs. 11.54 ± 1.91 for C2 of the contralateral hemicord) (Fig. 3C).

**Figure 3:**
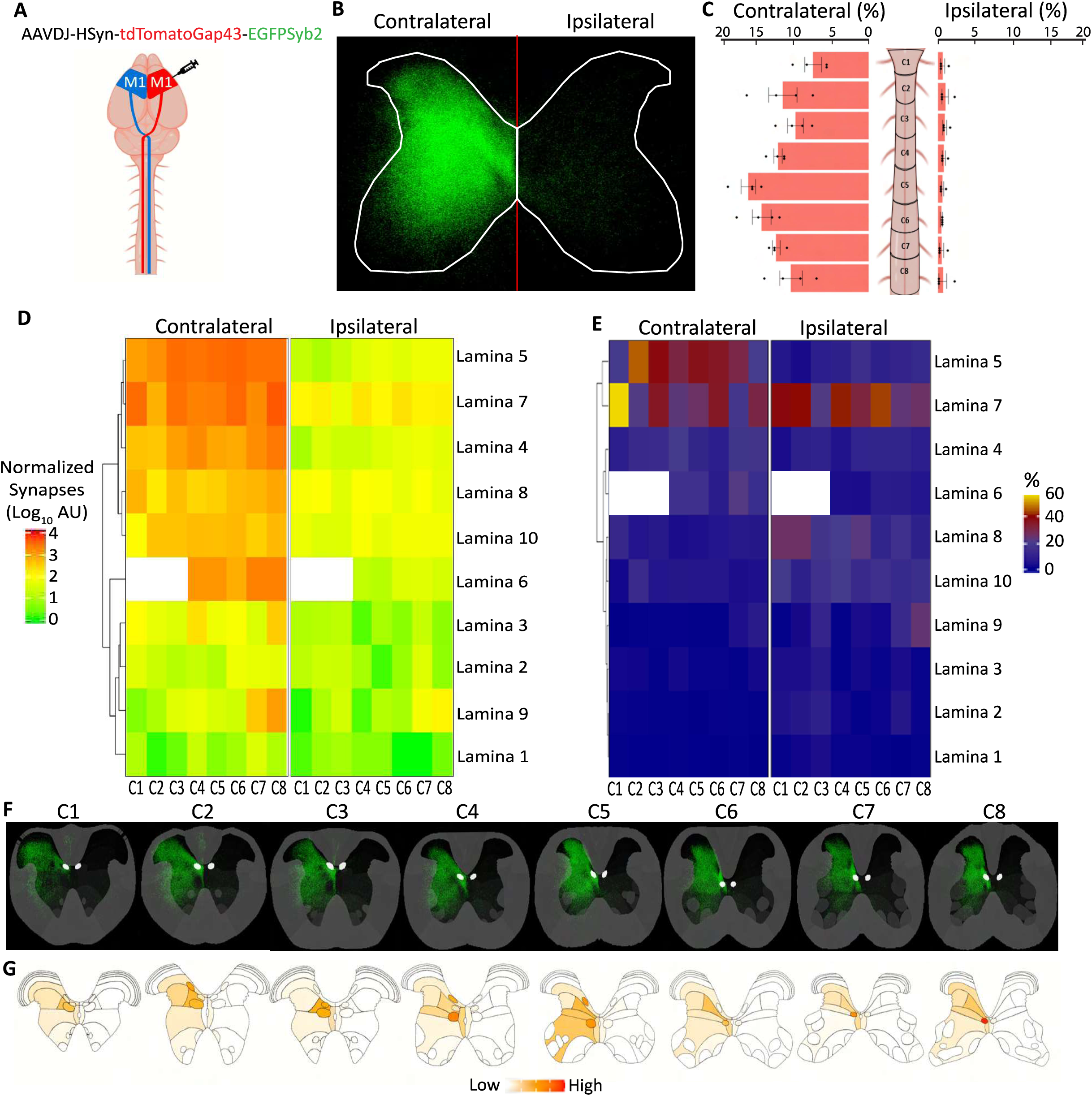
Region- and lamina-specific quantification of the cervical CST synaptic projectome in healthy adult mice. (A) Graphical representation of experimental design, showing a unilateral motor cortex injection of the AAV SynaptoTag4. (B) Combined probability maps of classified pre-synaptic terminal pixels across all images in the volumetric spinal cord dataset of a representative animal condensed and shown as a flattened image of the 3D volume. (C) Distribution of pre-synaptic terminals in each cervical level (C1-C8) of both the contralateral and ipsilateral hemicord, shown as the proportion (+/- SEM) of the total pre-synaptic terminals for each individual mouse. (D) Heatmap of pre-synaptic terminals showing the group mean of normalized synapses on a log10 scale at each cervical level and lamina. Lamina are organized according to dendrogram clustering based on pre-synaptic terminals in the contralateral hemicord. Empty boxes reflect that lamina 6 is not present at levels C1-3. (E) Distinct laminar distributions of pre-synaptic terminals in contra- and ipsilateral cords. Values represent relative percentage of the laminae for each cervical level (=100, vertical). (F) Pre-synaptic terminal probability maps for each cervical level (C1-C8) condensed and shown as flattened images of the 3D volumes. (G) Graphical depiction of mean synapse distribution in gray matter subregions (n = 4).

Examination of the lamina-specific distribution of synaptic terminals showed the highest density in laminae 5 and 7 in the contralateral hemicord (Fig. 3D, E; Supplemental Fig. 2).

Dendrogram analysis showed that laminae 5 and 7 clustered together in dendrogram analysis due to similarities in the total pre-synaptic terminals across most cervical levels (Fig. 3D) (Ueno et al., 2018). At most cervical levels, laminae 5 and 7 were the most densely innervated regions in the contralateral hemicord, containing up to 60% of pre-synaptic terminals within a given cervical level (Fig. 3D, E; Supplemental Fig. 2). Lamina 5 had nearly uniform presynaptic terminals across cervical levels C3-C8, and significantly less innervation of rostral cervical levels C1 and C2. In contrast, presynaptic terminals within Lamina 7 did not exhibit a rostral-to-caudal distribution pattern, with the densest innervation in C8, followed by C6 and C1 (Fig. 3D, F; Supplemental Fig. 2). Contralateral laminae 4 and 6 also clustered together in dendrogram analysis, showing a progressively caudal distribution of very modest innervation of cervical levels C4 and C5 and slightly higher innervation in C7 and C8 (Fig. 3D, F, G). Laminae 8 and 10 showed similar uniformly low presynaptic terminals throughout the caudal cervical levels (C4-C8). As expected, laminae 1-3 had very sparse pre-synaptic terminals throughout the entire contralateral hemicord of the cervical spinal cord, as reflected in the percent distribution of pre-synaptic terminals across laminae 1-3 for all cervical levels (Fig. 3D, E).

The ipsilateral hemicord exhibited a notably different distribution of CST pre-synaptic terminals across the laminae compared to the contralateral hemicord. Lamina 7 was a highly targeted region within the ipsilateral hemicord, with innervation ranging from approximately 20% to 40% at most cervical levels, mirroring the distributions seen in the contralateral hemicord (Fig. 3E). However, innervation of lamina 5 was markedly lower in the ipsilateral hemicord (2.64 to 14.36%) relative to the contralateral hemicord (18.88 to 45.89%) (Fig. 3E). Laminae 8 and 10 had the next highest terminal densities in the ipsilateral hemicord as opposed to laminae 4 and 6, which were favored in the contralateral hemicord at most cervical levels (Fig. 3D, F, G).

Overall, presynaptic terminals of laminae 4, 6, 8, and 10 in the ipsilateral hemicord was significantly higher and more rostral than in the contralateral hemicord (C1-C4: 29.9% vs 44.6%; C5-C8: 37.8% vs 43.1%; Contrateral vs Ipsilateral). In particular, in the ipsilateral hemicord, lamina 8 was heavily innervated, containing ∼25% of all pre-synaptic terminals at C1 and C2.

Furthermore, laminae 8 and 10 contained a greater percentage of the overall synapses at each cervical level in the ipsilateral hemicord relative to the contralateral hemicord (Fig. 3E).

### The CST directly connects to motoneurons in lamina 9 in healthy adult mice

We found that a proportion of the CST synaptic projections were located within motoneuron pools of lamina 9 in both the ipsilateral and contralateral hemicord (Fig. 3D). In fact, in the ipsilateral hemicord, Lamina 9 at cervical level C8 contained 23.8 ± 0.645% of the overall synapses at that cervical level and the presynaptic terminals in Lamina 9 at C8 was comparable to or greater than the presynaptic terminals of lamina 7 at many other cervical levels in that hemicord (C3, C7, & C8) (Fig. 3E). In line with this, raw fluorescent images clearly showed eGFP+ presynaptic terminals within the ventral horn in or around the motoneuron pools of Lamina 9 (Fig. 4A; Supplemental Fig. 3B). Maximum intensity projections of classified CST pre-synaptic terminals overlaid with ventral horn atlas images exclusively showing the motoneuron pools of caudal cervical levels, C6-C8, clearly demonstrated the abundance of these connections in both hemicords, albeit to a lower extent within the ipsilateral hemicord (Fig. 4B).

**Figure 4:**
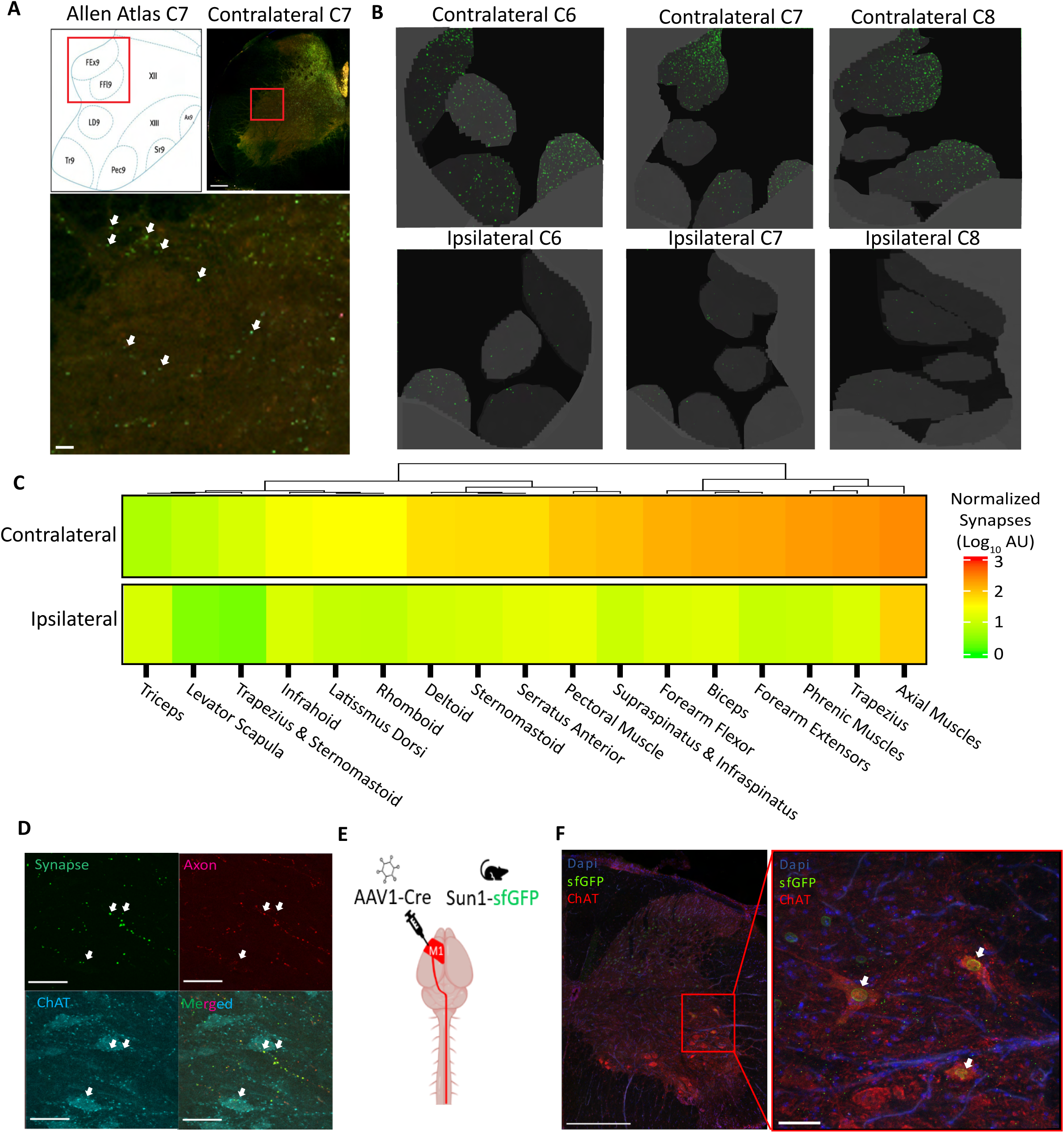
Direct CST synaptic connections with ChAT+ motoneurons in healthy adult mice. (A) eGFP+ pre-synaptic terminals in raw fluorescent image of a healthy adult mouse spinal cord section (scale bar 400um) with zoomed-in image of the ventral horn (red boxed area in upper right panel, scale bar 10um). White arrows denote pre-synaptic terminals. Upper left panel shows corresponding area of the atlas. (B) Cervical levels C6-C8 from the same animal in both hemicords showing condensed pre-synaptic terminal probability maps (green), masked to show only the ventral motoneuron pools (gray). (C) Quantification of mean pre-synaptic terminal levels for each motoneuron pool throughout the cervical cord across both hemicords, clustered using a dendrogram for the contralateral side. (N = 4) (D) Colocalization of synaptic terminals and ChAT+ motor neurons. Representative confocal z-stack of the contralateral ventral horn of a Synaptotag4-injected mouse immunostained with anti-ChAT antibodies. Synaptic GFP is shown in green, axonal tdTomato in red, and ChAT immunolabeling in blue (scale bar 50um). (E) Schematic representation of experimental setup for motor cortex injection of trans-synaptic AAV1-cre into the motor cortex of a Sun1-eGFP mouse. (F) Confocal images of a spinal cord section from an AAV1-injected Sun1-eGFP mouse, showing sfGFP+ nuclei (green) within ChAT+ motoneuron cell bodies (red) in a motoneuron pool within the contralateral hemicord. Left, whole hemicord image (scale bar 400um). Right, enlarged area boxed in red of a motoneuron pool (scale bar 50um).

Given the precisely characterized anatomical relationships of lamina 9 motor pools and their innervated musculature,(Stifani, 2014) we specifically assessed the distribution of CST pre-synaptic terminals across the cervical cord in the distinct motoneuron pools of lamina 9 based on the previously known innervated musculature. Similar to the other lamina, the distribution of CST presynaptic terminals within motoneuron pools differed between the ipsilateral and contralateral hemicord, although pre-synaptic terminals were found in every motoneuron pool in both hemicords (Fig. 4C, Supplemental Fig. 3A-B). The axial muscle motoneuron pool, uniquely present at all cervical levels, was bilaterally innervated by the CST synaptic projectome, albeit with more CST pre-synaptic terminals located in the axial muscle motoneuron pool in the contralateral hemicord relative to the ipsilateral hemicord (Fig. 4C). Innervation of the axial motoneuron pool was, by far, the most intensely targeted motoneuron pool in the ipsilateral hemicord, with very minor innervation of other motoneuron pools in the ipsilateral hemicord under homeostatic conditions. In contrast, many other motoneuron pools received substantial CST synaptic input in the contralateral hemicord, including the trapezius, phrenic muscle, forearm extensor, biceps, and forearm flexor motoneuron pools (Fig. 4C).

To identify whether motoneurons could be postsynaptic partners of the CST, spinal cord sections from SynaptoTag4-injected mice were immunostained with anti-ChAT antibodies, revealing eGFP+ CST pre-synaptic terminals are in close apposition with ChAT+ motoneurons within the contralateral lamina 9 (N = 3 mice, Fig. 4D). Next, we confirmed corticomotoneuronal monosynaptic connections using an AAV1 vector, AAV1-hSYN-Cre-P2A-dTomato (Zingg et al., 2017), which exhibits transsynaptic spread to first-order connections. Virus was injected into the primary motor cortex of CAG-SUN1/sfGFP mice (Mo et al., 2015), inducing Cre-dependent sfGFP expression in the nuclear membranes of postsynaptic neurons in the spinal cord (Fig. 4E-F) (Zingg et al., 2017; Zingg et al., 2020). We confirmed that sfGFP+ nuclei were present in all laminae previously identified with SpinalTRAQ as targets of the CST synaptic projectome (Supplemental Figure 3C), including lamina 9. Colocalization of ChAT immunoreactivity with AAV1-induced sfGFP+ nuclei in lamina 9 confirms that ChAT+ motoneurons receive direct synaptic input from the CST in the healthy, adult mouse (Fig. 4F, Supplemental Figure 3D).

### The post-stroke dynamics of the uninjured CST synaptic projectome

Recovery from primary motor cortex injury is associated with dramatic neuroplasticity in numerous pathways, including remodeling of the descending connections of the uninjured CST originating in the contralesional motor cortex (Carmichael et al., 2017; Chen et al., 2022; Liu et al., 2013; Liu et al., 2021). Around 4 weeks post-stroke, the midline-crossing collaterals of the uninjured CST increase in laminae 5-8 of the ipsilateral hemicord, which has been denervated due to stroke-induced CST degeneration (Kaiser et al., 2019; Liu et al., 2013). However, very little is known about the formation of new synapses by these sprouting axonal collaterals in the denervated hemicord, including the time course and anatomical location of synaptogenesis by the CST projectome. We used SpinalTRAQ to comprehensively assess the dynamic response of the uninjured CST spinal synaptic projectome to stroke, providing region-specific quantification of the temporal kinetics of spontaneous neuroplasticity during post-stroke functional recovery.

Adult C57/B6 mice received either a focal photothrombotic motor cortex stroke or sham surgery (Fig. 5A-B). We then labeled the contralesional, uninjured CST 2 weeks prior to euthanasia using viral SynaptoTag4 injection into the contralesional motor cortex and assessed post-stroke remodeling at 3 key recovery time points: 1-, 4-, and 6 weeks post-stroke (Fig. 5A) (Becker et al., 2016; Kaiser et al., 2019). At 1-week post-stroke, prior to the reported onset of CST axonal sprouting (Kaiser et al., 2019), there is very little difference in synaptic innervation of the denervated or ipsilateral hemicords of post-stroke and sham mice. (Fig. 5B, D, Supplemental Fig. 4A). No significant synaptic remodeling in any laminae was observed at this acute time point (Fig. 5D). However, by 4 weeks post-stroke, a time point where axonal sprouting is known to be maximal (Kaiser et al., 2019), a significantly greater percentage of CST synaptic terminals in the denervated hemicord were in C3 and C4 relative to sham animals (p<0.05; Fig. 5D) (Kaiser et al., 2019). At this time point, presynaptic terminals increased in the denervated laminae 5 and 7, particularly within lamina 7 at cervical level C4, mirroring healthy CST innervation patterns in the innervated hemicord of sham mice (Fig. 6A, Supplemental Fig. 4A). Additionally, increased presynaptic terminals were observed in laminae 4, 6, 8, and 10, regions containing interneurons that modulate descending CST motor signals, with the most notably elevated synaptic terminals in lamina 10 at cervical level C4 (Fig. 6A) (Gatto et al., 2021; Krotov et al., 2017; *The Mouse Nervous System*, 2011). Lamina 9 also showed modest increases in presynaptic terminals, suggesting the potential formation of corticomotoneuronal synapses at this time point (Fig. 6A; Supplemental Fig. 4A & B).

**Figure 5:**
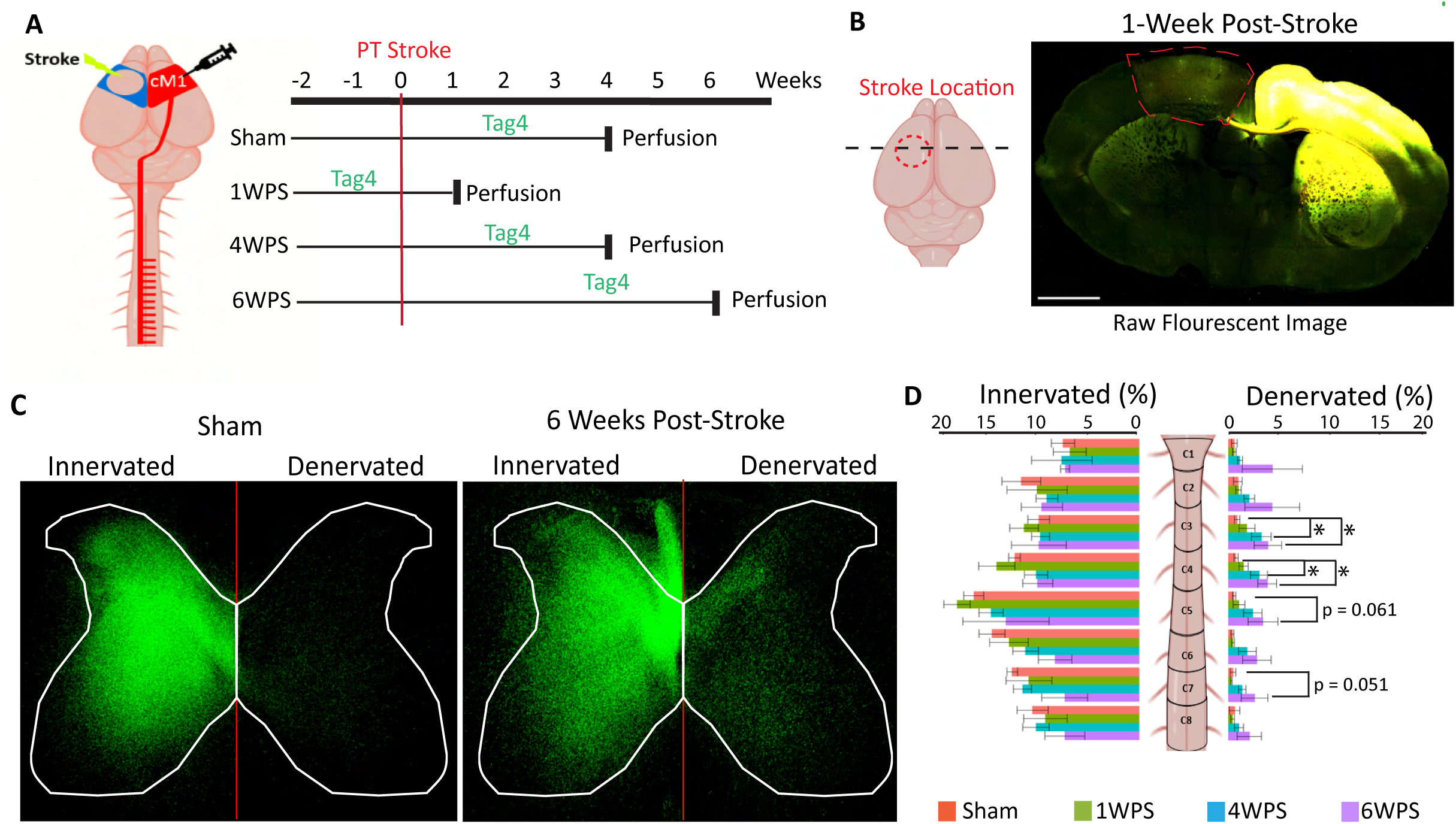
Post-stroke contralesional CST synaptic reinnervation in the denervated cervical hemicord. **(A)** Graphical diagram depicting experimental setup for photothrombotic (PT) stroke and injection of Synaptotag virus to contralateral motor cortex (scale bar 1mm). Mice were sacrificed at 1, 4, or 6 weeks post stroke (WPS). (B) Representative coronal image of stroke (traced in red) and virus injection area (yellow) at 1-week post-stroke imaged via STPT. (C) Comparison of condensed pre-synaptic terminal probability maps (C1-C8) from a sham and 6-week post stroke animal with similar pre-normalization viral labeling. (D) Distributions of CST pre-synaptic terminals at each cervical level (C1-C8) between both hemicords across all 4 groups, where each mouse’s entire spinal cord totals 100%, error bars are SEM. (Sham: n= 4, 1 WPS: n = 4, 4 WPS: n = 5, 6 WPS: n = 4)

**Figure 6:**
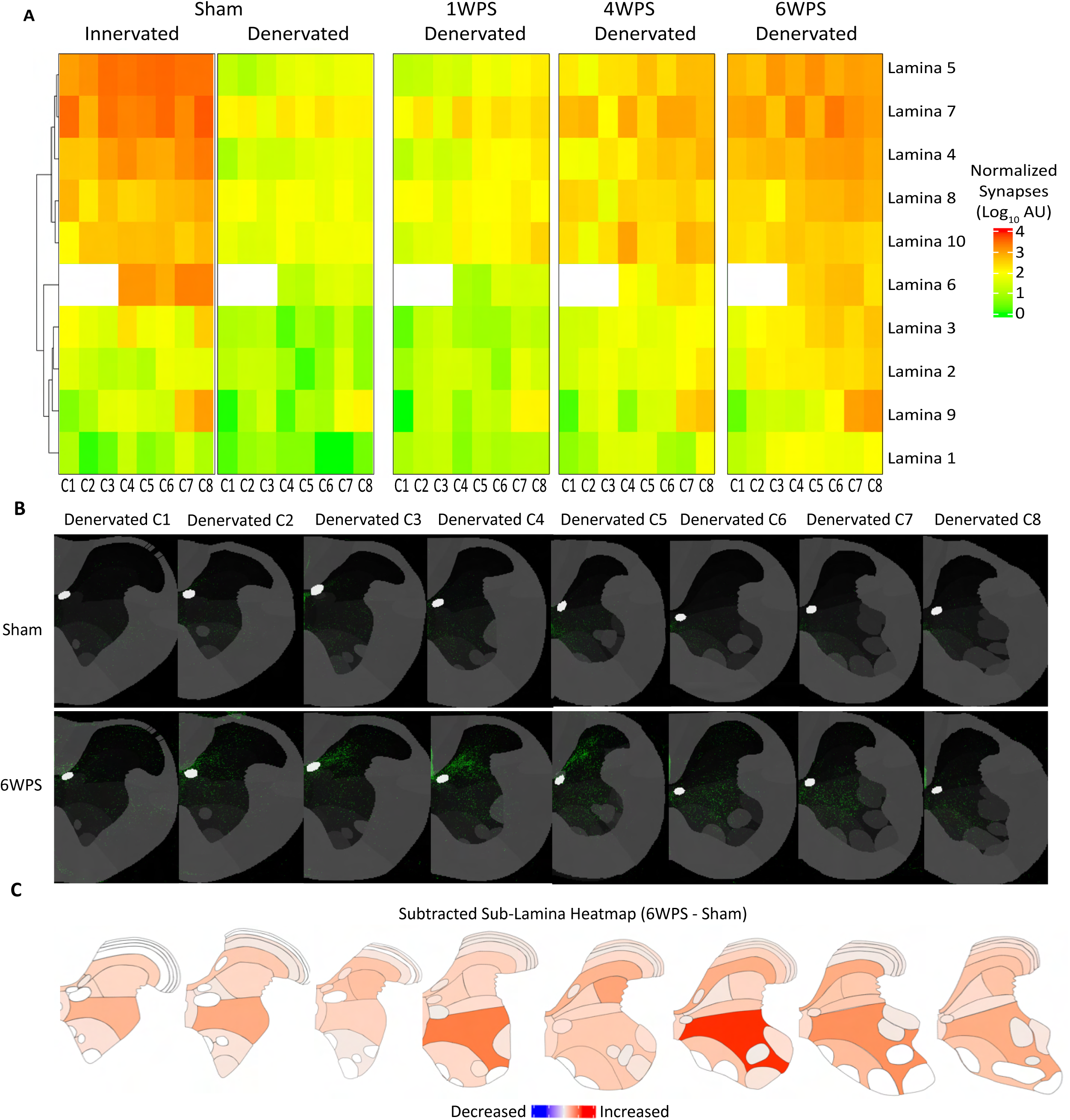
Level and lamina-specific reinnervation of the denervated hemicord by the contralesional CST synaptic projectome. (A) Heatmap demonstrating the mean of the group at each cervical level and lamina clustered using a dendrogram for the lamina of the sham innervated side. Only the denervated hemicord is shown for the three post-stroke groups. (B) Comparison of a sham and 6-week post stroke animal with similar pre-normalization viral labeling using condensed synapse probability maps across for each cervical level. (C) Heatmaps showing subtracted group mean of pre-synaptic terminals within atlas regions (6 weeks post stroke minus sham means). (Sham: n= 4, 1 WPS: n = 4, 4 WPS: n = 5, 6 WPS: n = 4)

At 6 weeks post-stroke, pre-synaptic terminals in the denervated hemicord increased dramatically (Fig. 5C-D, 6A, Supplemental Fig. 4A) with a significantly greater percentage of CST synaptic terminals located in C3 & C4 of the denervated hemicord compared to sham (p< 0.05; Fig. 5D). The CST synaptic projectome in the denervated hemicord of 6 weeks post-stroke mice demonstrated a strikingly similar laminar distribution to that of the contralateral CST in sham mice, with notable increases in presynaptic terminal density in lamina 7 at cervical levels C4 and C6 (Fig. 6A-C) Interestingly, despite its small overall size, lamina 4 also received robust synaptic innervation proportional to the reinnervation of laminae 5 and 7, perhaps due to reinnervation of the internal basilar nucleus, a region known to integrate descending input from the sensorimotor cortex with ascending sensory information (Fig. 6A, Supplementary Fig. 4a; (*The Mouse Nervous System*, 2011)). Modest innervation was observed in laminae 5, 7, and 10, again comparable to the contralateral innervation patterns seen in sham mice, with the disappearance of the increased innervation observed in lamina 10 at 4 weeks post-stroke (Fig. 6A-C).

### Post-stroke reinnervation of the denervated lamina 9 by the CST synaptic projection

In addition to the largely homotopic reinnervation of most laminae in the denervated hemicord, there is novel synaptic reinnervation of the denervated lamina 9 at caudal cervical levels beginning at 4 weeks post-stroke (Fig. 6A, Supplemental Fig. 4A). By 6 weeks post-stroke, synaptic innervation of all lamina 9 motor pools at cervical levels C7 and C8 exceeds innervation observed in lamina 5 and 7 at the same cervical levels (Fig. 6A-6C). The trend in caudal cervical levels towards a greater percentage of overall synapses in the denervated hemicord relative to sham mice (p<0.07; Fig. 5D) may be in part driven by the dramatic gains in presynaptic terminals across lamina 9.

Comparisons of the registered probability maps from the denervated caudal ventral horn show a visually appreciable increase in pixels classified as pre-synaptic terminals within motoneuron pools by 6 weeks post-stroke (Fig. 7A, Supplemental Fig. 4B). Further separation of lamina 9 into motoneuron pools based on the downstream innervated musculature reveals that in the ipsilateral hemicord of sham mice, nearly all motoneuron pools receive minimal CST synaptic input with the exception of the axial muscle motoneuron pool (Fig. 7A). At 1 week post-stroke, similar to other laminae, there is minimal reinnervation of any denervated lamina 9 motoneuron pools (Fig. 7B). By 4 weeks, there is denser innervation of the denervated axial muscle motoneuron pools relative to the ipsilateral axial muscle motoneuron pool (Fig. 7B). At this time point, innervation also increases in additional motoneuron pools, including the forearm flexor (Fig. 7B).

**Figure 7:**
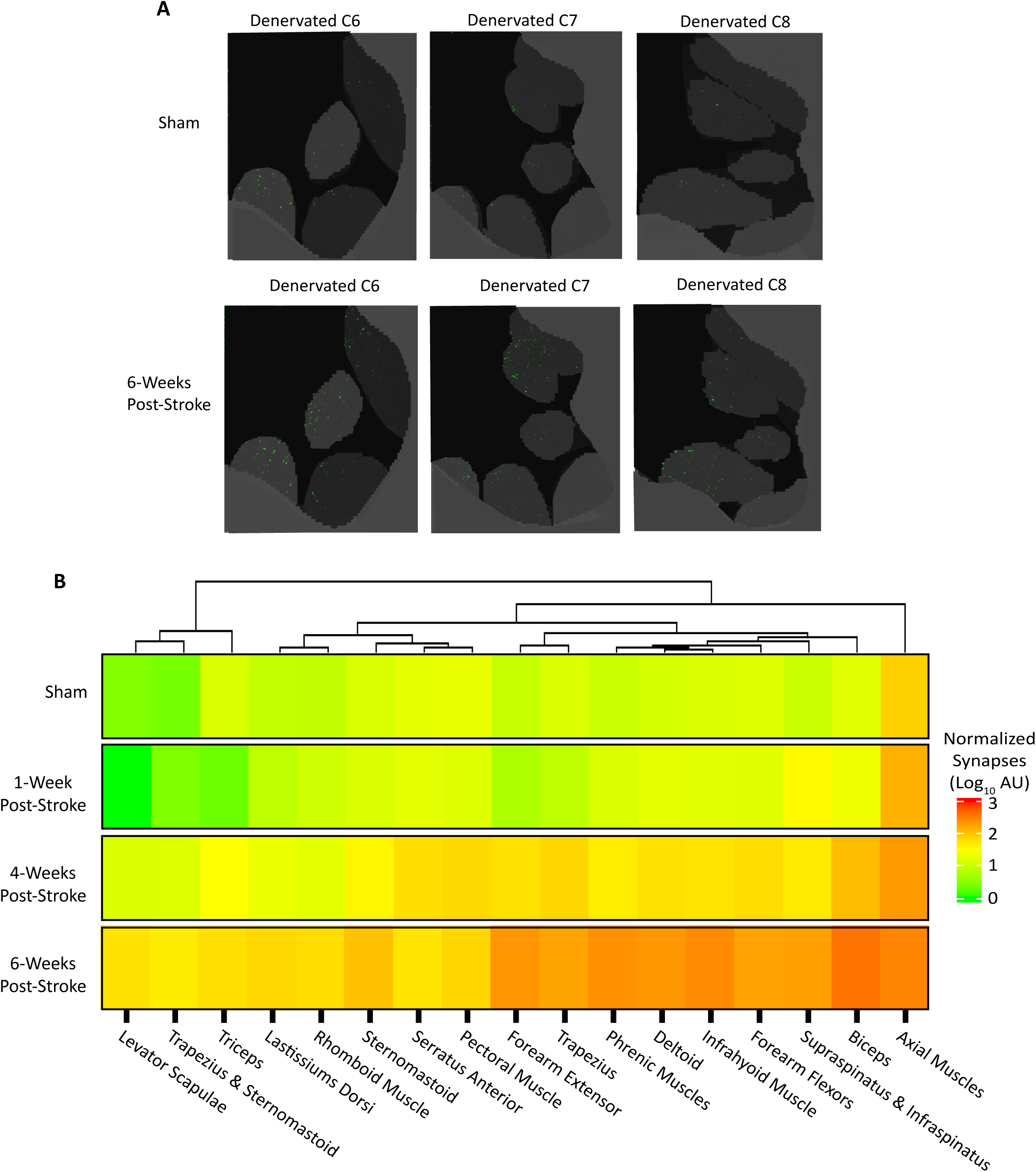
Increase in direct CST synaptic connections within specific laminae 9 motor pools after stroke. (A) Cervical levels C6-C8 from a sham and 6-week post stroke animal with similar pre-normalization viral labeling showing condensed pre-synaptic terminal probability maps for each cervical level. Only ventral motoneuron pools are shown for clarity. (B) Heatmap demonstrating the mean of the group at each laminae 9 motor pool of the denervated hemicord annotated in the atlas, clustered using a dendrogram for sham denervated. (Sham: n= 4, 1 WPS 1: n = 4, 4 WPS: n = 5, 6 WPS: n = 4)

By 6 weeks post-stroke, there is a notable increase in CST synaptic reinnervation of the denervated lamina 9 (Fig. 6A) in nearly all motoneuron pools (Fig. 7A, Supplemental Fig. 4B). In contrast to the global reinnervation of denervated premotor laminae at 6 weeks that occurs throughout the entire cervical cord, (Fig. 6A) post-stroke reinnervation of motoneuron pools exhibits a strong bias in the rostral-caudal axis, heavily favoring reinnervation in the caudal cervical levels, C7 and C8 (Fig. 6A). Furthermore, unlike reinnervation of premotor laminae, post-stroke CST projections to denervated motoneuron pools is not homotopic (Fig 7A). In the contralateral hemicord of healthy adult male mice, the forearm flexor receives only moderate CST synaptic input relative to the more innervated axial muscles (Fig. 4C). However, in the denervated hemicord of 6 weeks post-stroke mice, the forearm flexor is the most densely innervated motoneuron pool, demonstrating novel post-stroke patterns of potential corticomotoneuronal connectivity. Thus, these SpinalTRAQ results reveal unprecedented detail about reinnervation of the denervated hemicord at the heretofore undescribed synapse level, showing both homotopic reinnervation of premotor laminae as well as novel connectivity within caudal motoneuron pools in lamina 9.

## Discussion

To our knowledge, this is the first unbiased, quantitative bilateral measure of the CST synaptic projectome, localizing pre-synaptic terminals with an unprecedented degree of level and region specificity across the entire cervical cord volume. Building on a recently reported digital atlas of the mouse spinal cord (Fiederling et al., 2021a, 2021b), we describe specific topography of CST synapses in health and after stroke. SpinalTRAQ has several advantages over conventional serial sectioning and immunofluorescent staining methods (Liu et al., 2013): it eliminates time-consuming cryosectioning and immunofluorescent staining, reduces sampling bias, and minimizes tissue distortion and artifacts. We validate our atlas classifications using transgenic ChAT-GFP mice and successfully detail the homeostatic and remodeled CST spinal synaptic projectome using this novel approach. The platform can be readily adapted to assess many cervical spinal cord features with anatomical region specificity. However, we also acknowledge that SpinalTRAQ in its current form has limitations, including that STPT requires the presence of an endogenous fluorescent signal, with very few options for exogenous labeling of structures of interest (Poinsatte et al., 2019; Ramirez D. M. O., 2019). Future work will adapt SpinalTRAQ to be compatible with other popular high-resolution volumetric imaging techniques such as light sheet microscopy, further expanding the accessibility and utility of the method.

In this study, we utilize SpinalTRAQ to generate the CST synaptic projectome in the mouse cervical spinal cord. The precise region specificity of SpinalTRAQ not only reveals the dense largely unilateral synaptic innervation of laminae 5 and 7 in the contralateral hemicord, regions previously shown to be rich in premotor interneurons and major targets of CST axonal projections (Bui et al., 2013; Levine et al., 2014), but also highlights the sometimes overlooked sparser CST synaptic connections in the ipsilateral hemicord of healthy, adult mice. The CST synaptic projectome in ipsilateral hemicord exhibits a unique innervation pattern relative to the more highly innervated contralateral hemicord. Most of the CST pre-synaptic terminals in the ipsilateral hemicord of the healthy adult mouse are found prominently within lamina 7 and more modestly within laminae 8 and 10, but not within laminae 4 or 5, regions that receive comparatively more CST synaptic input in the contralateral hemicord. Our findings of CST synaptic innervation in the ipsilateral hemicord are consistent with reports of axonal innervation of the ipsilateral hemicord by midline-crossing CST collaterals pre-injury in control healthy adult mice (Kaiser et al., 2019; Lapash Daniels et al., 2009; Ueno et al., 2012). While midline-crossing collaterals could be targeting errors, as research using EphA4/EphA3 knockouts have suggested (Chedotal, 2019; Kullander et al., 2001; Paixao et al., 2013; Restrepo et al., 2011), we hypothesize that the unique innervation pattern in the ipsilateral hemicord reflects a discrete, currently unknown biological function for these CST synaptic connections in laminae 7, 8, and 10. Presented here is an improved understanding of the distinct topography of these sparser connections in the ipsilateral hemicord of healthy mice, laying the groundwork for future investigation into the physiological roles of these CST connections in the modulation of spinal neuron populations.

In addition to detailing the nuanced innervation of laminae containing premotor interneurons in the ipsilateral hemicord, SpinalTRAQ has further revealed bilateral corticomotoneuronal synapses in lamina 9 of healthy adult male mice, particularly within caudal motoneuron pools, including those controlling forelimb muscles. Direct synaptic input from the CST onto motoneurons is believed to underpin the increased manual dexterity of primates (Bortoff & Strick, 1993; Heffner & Masterton, 1975). Primate studies have shown that the number of corticomotoneuronal synapses into forelimb motoneuron pools directly correlates with increased hand grip strength (Armand et al., 1997; Olivier et al., 1997). While these corticomotoneuronal connections exist during development for mice (Gu et al., 2017), evidence for direct synaptic input from the CST onto motoneurons in healthy adult mice is limited (D’Acunzo et al., 2014; Kuang & Kalil, 1990) and controversial (Lemon, 2019; Lemon, 2008).

Therefore, we confirmed our unbiased machine learning-driven classification of pixels as CST pre-synaptic terminals within caudal lamina 9 motoneuron pools with additional transsynaptic viral tracing experiments, in which AAV1-Cre induced sfGFP expression in the nuclei of spinal cord neurons that directly received input from the CST synaptic projectome. Clear colocalization of ChAT immunostaining with sfGFP+ nuclei in the ventral horn confirms the direct monosynaptic nature of the corticomotoneuronal connections. Because AAV1-mediated labeling of post-synaptically connected neurons requires the transmission of viral particles across functional synapses with active synaptic release, this implies that observed corticomotoneuronal synapses may have functional relevance to mouse forepaw dexterity (Zingg et al., 2017; Zingg et al., 2020). Supporting these results, a recent publication using a similar AAV1 transsynaptic viral labeling technique to characterize the spinal postsynaptic partners of the CST found approximately 6% of the spinal target neurons were ChAT+, but did not specifically delineate the anatomical location of these ChAT+ postsynaptic neurons (Carmona et al., 2024). The comprehensive laminae- and level-specific analysis of SpinalTRAQ uniquely allowed us to identify the sparse, but potentially critical, corticomotoneuronal contacts, highlighting the unique spatial sensitivity of our methods. However, a trade-off of the anatomical rigor of our SpinalTRAQ methodology is the potential slight undercounting of corticomotoneuronal synapses. In our pixel classification of the GFP+ motoneuron dendrites in ChAT-GFP reporter mice, we found a percentage of these dendrites projected far outside of lamina 9 into laminae 7 and 8. Likely corticomotoneuronal synapses on motoneuron dendrites in laminae 7 or 8 could be overlooked, indicating the limitations of an atlas based on predetermined atlas regions rather than exclusively connectivity.

After stroke, dynamic changes in the CST synaptic projectome occur throughout the ipsilateral denervated hemicord. These can be broadly divided into two types of post-stroke CST synaptic projectome remodeling: (1) homotopic reinnervation of premotor laminae and (2) novel connections within lamina 9. Beginning 4 weeks after a motor cortex stroke, synaptic terminals increase not only in lamina 7, but also within laminae 4 and 5, in the denervated hemicord. This suggests that the sprouting CST engages in a homotopic reinnervation of laminae 4, 5, and 7 interneuron populations, such as V2a (Hayashi et al., 2018), motor synergy encoders (Levine et al., 2014), and dl3 premotor interneurons (Bui et al., 2013) that have lost CST synaptic input due to stroke-induced CST degeneration and are known to contribute to recovery after injury (Kathe et al., 2022; Kumamaru et al., 2019). In rats, the precise targeting of particular anatomical regions within the denervated hemicord by sprouting CST axons post-stroke, particularly laminae 6-9, has been shown to be required for motor recovery, as therapies promoting excessive plasticity with aberrant laminar fiber patterns favoring dorsal laminae did not confer the same fine motor skill benefits (Wahl et al., 2014). We hypothesize that CST synaptic remodeling in mice must similarly occur in a topographically controlled, mostly homotopic manner. However, more studies are required to establish a link between anatomically sensitive synaptic remodeling and motor function.

In addition to homotopic reinnervation of premotor laminae, we observed unique changes in CST connectivity within lamina 9 after stroke. Alterations in the CST synaptic projectome within lamina 9 differed from homotopic reinnervation of premotor laminae in two prominent ways. First, while the bilateral innervation of motoneuron pools by the CST synaptic projectome is relatively minor in healthy mice, CST input to the caudal denervated lamina 9 motoneuron pools markedly increases by 6 weeks post-stroke, reaching densities comparable to or exceeding that seen in laminae 4, 5, and 7 at many cervical levels. Therefore, in addition to a likely reestablishment of prior patterns of homeostatic connectivity in laminae 4, 5, and 7 at 4 weeks post-stroke, the increase in corticomotoneuronal synapses at 6 weeks post-stroke may comprise a novel pattern of CST synaptic innervation not normally observed in healthy adult mice. In line with this hypothesis, of the few studies that suggest direct CST input to motoneurons, several come from spinal cord injury research (Steward et al., 2004; Steward et al., 2008). However, we are the first to link corticomotoneuronal connectivity to more remote injuries such a motor cortex stroke. Second, while the likely homotopic reinnervation of premotor interneurons occurs somewhat uniformly throughout the cervical spinal cord, we observed that increased corticomotoneuronal synapses in the denervated lamina 9 differed more strikingly across the rostral-caudal axis of the spinal cord. Within the denervated lamina 9, nearly all changes in connectivity occurred in caudal cervical levels C7 and C8, the levels in which forelimb flexor and extensor-controlling motoneurons reside. In line with this, we observed increased innervation by the CST synaptic projectome within the forearm flexor-innervating motoneuron pool. Because our PT stroke model specifically targets forelimb-related regions of the motor cortex and induces long-lasting impairments in forelimb function requiring precision and dexterity (Becker et al., 2020; Becker et al., 2016), the increase in pre-synaptic terminals to levels exceeding homeostatic connectivity, particularly in caudal forelimb motoneuron pools, may reflect plasticity aimed at ameliorating forelimb-specific dexterity deficits after stroke.

SpinalTRAQ permits unbiased lamina- and level-specific quantification of spinal cord phenomena in 3D volumetric space utilizing a novel workflow which incorporates serial two-photon tomography and automated registration to a custom cervical cord atlas. Amongst the many potential broad applications of this method, we demonstrate how it can illuminate the finer details of CST connectivity under homeostatic conditions, including strong evidence for minor, but potentially meaningful, connections with motoneurons. Furthermore, SpinalTRAQ reveals the time course of dynamic changes in the CST synaptic spinal projectome over the course of stroke recovery, highlighting unique temporal and spatial dynamics of both broad homotopic reinnervation of laminae 4, 5, and 7 and the injury-specific corticomotoneuronal connectivity in caudal lamina 9. Future studies should explore the broad potential applications of this methodology as well as delve deeper into the functional relevance and implications of the spatially-specific CST synaptic spinal projectome.

## Materials & Methods

### Animals

Experiments were performed in accordance with protocols approved by either UT Southwestern’s Institutional Animal Care and Use Committee (IACUC) or UT Health Science Center San Antonio’s IACUC. All CST synaptic projectome experiments were performed on 8-11 week old male C57BL/6 mice (n=30) from Jackson Laboratories. ChAT-GFP experiments were performed on 9 week old transgenic mice (Jax # 006410 or # 007902) (n=3) expressing eGFP in cholinergic ChAT+ neurons throughout the central nervous system. For AAV1-Cre tracing experiments, male 8-16 week old Sun1-sfGFP (Jax # 021039) mice (n=3) were used. Animals were housed in a 12 h light/dark cycle with access to food and water ad libitum.

### Photothrombotic (PT) Stroke

Motor cortex stroke was induced using the photothrombotic stroke model previously described (Yanev et al., 2020). Mice (n = 6) were anesthetized using 1–4% isoflurane, in a mix of 70% nitrous oxide and 30% oxygen, and temperature and breathing rates were monitored. Mice were placed on a stereotaxic frame and an incision was made down the midline of the scalp utilizing aseptic technique. Mice were administered Rose Bengal (Sigma Aldrich, St.

Louis, MO, USA; 40 mg/kg, dissolved in saline) via intraperitoneal injection. One minute later, a 45 mW laser (Coherent Sapphire, Santa Clara, CA, USA; 561 nm; 2.7 mm collimated beam diameter) was activated for 15 min (aimed 1.7 mm lateral to Bregma to target the forelimb representation of the left motor cortex). Sham mice underwent the same procedure outlined above (anesthesia, Rose Bengal, stereotaxic mounting, etc.) for the same amount of time but did not have laser exposure. aBuprenorphine ER was administered post-operation for pain management, and moist food was provided for the first 24-72 hours following stroke. All mice that received a PT stroke had successful surgeries. PT strokes were performed by the Neuromodels Facility at UT Southwestern Medical Center (RRID:SCR_022529).

### Virus production of SynaptoTag4

SynaptoTag2 was modified from SynaptoTag4 to include a GAP43 promoter to target tdTomato expression to axonal membranes and packaged into an AAVDJ serotype. It was prepared as described in (Li et al., 2021). Briefly, AAV vectors were cotransfected with pHelper and pRC-DJ into AAV-293 cells. 72 hours later, cells were collected, lysed, and centrifuged at 400,000g for 2 hours in an iodixanol gradient. The 40% iodixanol gradient fraction was washed and concentrated with a 100,000 molecular weight cutoff tube filter. The titer of SynaptoTag4 used was within the range of 0.5-2 x 10^13^ genome copies per mL, as determined by quantitative real-time PCR. Virus was aliquoted and frozen at -80C until use. SynaptoTag4 virus was generously provided by Wei Xu’s laboratory at UT Southwestern.

### Intracortical AAV injections

For CST synaptic projectome experiments, mice were anesthetized with 1–4% isoflurane, in a mix of 70% nitrous oxide and 30% oxygen and temperature and breathing rates were monitored. Mice were placed on a stereotaxic frame, and following fur removal and skin preparation utilizing aseptic technique an incision was made down the midline of the scalp. A small burr hole was drilled in the skull at 1.5 mm lateral to Bregma, in accordance with the forelimb representation of the motor cortex. Virus was loaded into a 2.5 uL Hamilton syringe (Hamilton #7535-01) backfilled with neutral mineral oil and fitted with a pulled glass pipette using an RN Compression Fitting (Hamilton #55750-01). The glass pipette was lowered to a depth of -0.6mm and a pressure injection of 1 uL of AAV (SynaptoTag4) was made over 10 minutes into the right motor cortex, followed by a 10-minute waiting period before the glass pipette was removed. For AAV1-cre injections, Sun1-GFP mice were anesthetized using 1-2% isoflurane and oxygen and underwent scalp shaving, topical lidocaine, and an incision down the midline of the scalp. Following cleaning of the skull with hydrogen peroxide to reveal bregma, mice were mounted to a custom software controlled motorized stereotaxic instrument and received a single unilateral injection via a pulled glass micropipette of AAV1-cre of either 1ul (n=2) or 500nl (n=2) at -0.2/1.5 (A/P relative to bregma) and -0.5mm below cortical surface into the right motor cortex at a rate of 100nl/min. The pipette was left in place for 1 minute and then raised 0.2mm, followed by an additional 1 minute static hold and then subsequently removed. pAAV-hSyn-Cre-P2A-dTomato was a gift from Rylan Larsen (Addgene viral prep # 107738-AAV1; 2x10^13^). AAV1-Cre induces GFP expression in postsynaptically connected neurons via transmission of viral particles across functional synapses with active synaptic release, in transgenic mice with Cre-dependent expression of GFP (Zingg et al., 2017; Zingg et al., 2020). Buprenorphine was administered post-operation for pain management, and moist food was provided for the first 24-72 hours following stroke.

### Sample preparation for imaging on TissueCyte 1000

For CST synaptic projectome experiments, all mice were sacrificed 2 weeks after Synaptotag4 injections via transcardial perfusion with 1X Phosphate Buffered Saline (PBS) followed by 4% paraformaldehyde (PFA) to fix brain and spinal cord tissues. Extracted tissues were further post-fixed in 4% PFA at 4◦C for 24 h, then the cervical segment of the spinal cord was separated from the brain and thoracic cord and transferred to 0.01% NaN3 in 1X PBS for storage at 4◦C until embedding. 4% Surecast solution (Acrylamide:Bis-acrylamide ratio of 29:1, ThermoFisher #HC2040) with 0.5% VA-044 activator (Wako, Catalog #: 27776-21-2) was prepared on ice, and cervical spinal cord segments were soaked in this solution overnight at 4◦C before being removed and placed at a shaker at room temperature for 1 hour to equilibrate. The cervical spinal cord segments were transferred to a disposable mold (VWR, Catalog #: 15160-215), and excess acrylamide solution was added (2.5ml) to cover the tissues completely. Molds were covered tightly with aluminum foil and placed in an oven preheated to 40◦C for 2 to 3 hours. Simultaneously, 4.5% (w/v) agarose (Type 1A, low EEO, Sigma #A0169) solution in 50 mM phosphate buffer was prepared and oxidized by adding 10 mM NaIO4 (Sigma #S1878) while stirring gently for 2 to 3 h in the dark. The agarose solution was filtered with vacuum suction and washed 3 times with 50 mM phosphate buffer (PB). The washed agarose was resuspended in the appropriate volume of PB to make a 4.5% (w/v) agarose solution. 10ml of agarose solution is needed for each spinal cord segment to be embedded. The oxidized agarose solution was heated to boiling in a microwave, then transferred to a stirring plate and allowed to cool to 60–65◦C. Spinal cord segments were removed from the oven and molds and excess agarose was gently removed from the tissues using Kimwipes or tweezers. The cervical segments were embedded by placing a fresh cryoembedding mold (VWR #15560-215) on a flat frozen ice pack, partially filling with 3ml oxidized agarose heated to 60-65◦C, and then quickly submerging the cervical sections using forceps into the bottom of the block with the rostral side touching the bottom of the block (caudal side facing upward). Once the agarose had hardened slightly around the tissue, the rest of the block was filled with agarose until slightly overfilled using a disposable transfer pipette. The agarose block was allowed to fully solidify on a frozen ice pack in the dark. Once the agarose blocks were fully hardened, the tops of the agarose blocks were trimmed with a razor blade until flat and level with the sides of the mold, then the blocks including the specimens were removed from the molds and placed in individual small glass jars (Uline #S-17073M-W), where they were treated overnight at 4◦C in the dark in sodium borohydride buffer (50 mM sodium borohydride (Sigma # 452882), 50 mM borax (Sigma, #221732), 50 mM boric acid (Sigma # B6768), pH 9.0–9.5). After overnight crosslinking, the agarose blocks were transferred to phosphate buffer for storage at 4◦C until TissueCyte imaging.

### Serial two-photon tomography (STPT)

The agarose blocks containing the spinal cord samples were attached to a custom magnetic slide with superglue and placed on a magnetized stage within an imaging chamber filled with phosphate buffer. STPT imaging is a block-face imaging technique in which a series of 2-dimensional (2D) mosaic images in the transverse plane are acquired just below the cut surface of the spinal cord, followed by physical sectioning with a built-in vibrating microtome to cut away the imaged tissue, preparing a new cut surface for imaging (Ragan et al., 2012). For this study, three optical planes were imaged at 20, 40, and 60 μm below the cut surface, followed by a vibrating microtome cut at 60 μm (blade vibration frequency of 60 Hz, advancement velocity 0.5 mm/s). The excitation laser (MaiTai DeepSee, SpectraPhysics/Newport, Santa Clara, CA) wavelength was tuned to 780 nm for initial sample positioning, then 930 nm for image acquisition to efficiently excite both GFP and tdTomato present in the cord segments. Spinal cords were positioned for imaging by centering the central canal beneath the objective before moving the stage to the starting position. Three emission channels using pre-set bandpass filters encompassing red, green, and blue emission were collected with a predetermined photomultiplier tube voltage of 720 V. This process produced 57,600 image tiles, 200 physical sections, and 1,800 2D stitched coronal section images (8 by 12 mosaic, 1 stitched coronal section image for each channel and each optical plane) with a lateral resolution of 0.875 μm/pixel and axial resolution of 20 μm (∼280 gigabytes of raw data per cord). The raw image tiles were first trimmed and subjected to flat field correction, then stitched into 2D mosaic coronal section images using the Autostitcher software (TissueVision, Inc.). Single channel, single plane stitched coronal section images were deposited on our cluster computing resource, BioHPC, for further processing and analysis using our custom developed pipelines. TissueCyte imaging was performed in the UT Southwestern Whole Brain Microscopy Facility, RRID: SCR_017949.

### Image Analysis Pipeline

We used an automated pipeline (similar to the whole mouse brain analysis pipeline described in (Poinsatte et al., 2019; Ramirez et al., 2019) to extract quantitative results from raw STPT-generated cervical spinal cord data. The image analysis pipeline consists of three main components: image registration, pixel classification, and numerical quantification. We have created a novel 3D mouse cervical spinal cord atlas to enable this process, which consists of a custom STPT based cervical spinal cord average template volume, and the digitized anatomical region annotations available as part of the SpinalJ analysis pipeline (Fiederling et al., 2021a, 2021b).

The raw fluorescence signal from the red channel of the stitched 3D images is used to register the dataset to our STPT based cervical spinal cord average template using a custom pipeline constructed with the Simple Elastix package (Lowekamp B., 2015). Cord samples were padded with blank section images to 200 sections in the Z-dimension if fewer than 200 sections were collected during imaging or cropped to 200 Z sections if more than 200 sections were collected, to ensure all cords had the same image dimensions for registration. Because curvature along the length of the spinal cord in the Z-dimension can cause the section images to be horizontally and vertically displaced from the center of the atlas template, section images were automatically re-centered using a MATLAB script during pre-processing to improve the registration initialization. A multi-step registration process was chosen to use rigid, affine, and b-spline registration components. At each step, optimization was performed on a 6-resolution multi-grid (pyramid) schedule, with Mattes Mutal Information as the optimization metric. Final b-spline grid dimensions were 1 x 1 x 1 mm. In cases where the automated registration alignment was not optimal, an additional registration step was performed, in which the Euclidean distance between manually specified fiducial markers in the sample and atlas was minimized (see below for further details). The optimized registration transformation identified for the raw red channel images was then applied to the other raw image channels.

In parallel to identification of registration parameters for each sample, relevant signals of interest were identified in the images using the “Pixel Classification” plugin for the Interactive Learning and Segmentation Toolkit (ilastik), which implements a random forest supervised machine learning (ML) classifier (Somner, 2011). To support the supervised ilastik training, a maximum intensity projection (MIP) was first created for each physical section from its three optical planes. The resultant MIP image was color adjusted to improve fluorescent signal contrast, and downsampled to a pixel size of 1.5 μm × 1.5 μm in the x-y dimension. This resulted in a set of 16-bit 3-color compressed TIFF images with the same number of images as physical sections in the original STPT image volume. Up to five representative sections containing examples of fluorescent signals of interest (eGFP+ pre-synaptic terminals/tdTomato positive axons or eGFP+ cholinergic neurons) and spanning the rostro-caudal extent of the cervical spinal cord sample were selected from each sample. The random forest ML model was trained on these selected sections by labeling pixels as axons, pre-synaptic terminals, white matter, gray matter, and other features (noise, background, etc). After completing the initial training, each of the sample images was iteratively updated until visible misclassifications were minimized. Each cohort of spinal cords (CST synaptic projectome or ChAT-GFP validation experiments) was used to train an independent model for classification, and every section from every spinal cord in the cohort was subjected to the same model. Ilastik predictions were exported as an 8-bit “probability map” TIFF image for each trained label, with the pixel value of 0 mapping to 0% probability and 255 100% probability of a pixel belonging to the label. The resulting TIFF stacks were warped to the custom STPT-based cervical spinal cord average template using the transformation parameters identified during the registration process for the raw fluorescent images. The probability maps were then thresholded (pixel threshold = 86, determined visually) to remove low probability noise. The summed intensity of all denoised probability map voxels lying within annotated regions present in the atlas were quantified using custom MATLAB software, producing a matrix of signal intensity for each sample, label, and cervical spinal cord region.

### Manual Fiduciary Point Placement

In cases where raw image autofluorescence differed substantially from the average template, manual fiduciary point placement was used to fix misaligned images within individual animals. The average template and autofluorescence channel were both opened in ImageJ, and the point tool was used to add points on both image sets that correspond to tissue architecture in both image sets. Routinely marked regions include central canal, gray/white matter boundary with the dorsal funiculus at the midline, gray/white matter boundary of the ventral funiculus at the midline, lateral and medial edges of the dorsal horns, ventral most point of the gray/white matter boundary, and substantially complex ventral horn gray/white matter boundary of lower cervical level such as the forearm motor pool. On average 50-100 points are selected in a given sample. A custom ImageJ script was used to obtain coordinates of both images with the point indices.

### Immunofluorescence

For validation of corticomotoneuronal synapses in AAV1-Cre injected mice, mice were transcardially perfused with 1X PBS followed by 4% PFA. Cervical spinal cords were dissected and fixed overnight in 4% PFA, followed by 2 nights of incubation in 30% sucrose in 1X PBS (w/v). Spinal cords were sectioned via microtome at 45um, photobleached utilizing a X-cite120 metal halide lamp and a Quad transmission/excitation filter (Chroma: 89401: 405/488/555/640nm) for 5 minutes, and stained with anti-ChAT primary antibodies (EMD Millipore Sigma:AB144P, 1:200; Thermofisher:CL3173 1:200) and anti-GFP (Thermofisher: G10362, 1:200) with secondary antibody donkey anti-goat Alexa488 (Thermofisher: A11055; 1:500) and donkey anti-rabbit Alexa564 (Thermofisher: A10042; 1:500) or goat anti-mouse Alexa647 (Thermofisher: A21236; 1:500). Sections were mounted, cover slipped, and images were acquired using a Zeiss LSM710 confocal microscope at 25x magnification.

### Assessment of viral labeling efficiency

Thoracic and lumbar spinal cords were collected from Synaptotag injected mice at the time of sacrifice, stored for 24 hours in 4% PFA and then cryoprotected in 30% sucrose for a minimum of 2 nights. Thoracic and lumbar spinal cords were embedded in OCT and cryosectioned at 30-40 um. After DAPI staining (1:5000), sections were mounted, coverslipped and whole slide images were acquired on the Zeiss Axioscan.Z1 in the UT Southwestern Whole Brain Microscopy Facility (RRID:SCR_017949). Images (data not shown) were assessed to determine if sufficient viral labeling of the CST was achieved. Robust CST labeling was observed in 21 animals, and their cervical spinal cords were processed for STPT and SpinalTRAQ analysis. Nine animals were excluded because the viral labeling observed was insufficient for further analysis.

### Statistics

Data were analyzed in R (R version 4.1.2; The R Foundation for Statistical Computing) using custom scripts. Synapse quantification was normalized to control for injection efficacy and spread by dividing each animal by its left hemicord classified axon intensity to cover CST regions outside of the dorsal column. SpinalTRAQ produces two different quantifications, raw intensity, a measure of total positive pixels, and density of positive pixels (raw intensity/volume of atlas region = density) for every atlas region. We chose to report normalized raw intensity values in main figures in effort to not require the reader to have biological context of the volume of all spinal cord regions across all 8 cervical levels. In addition, the quantification of CST synapses in a density manner currently lacks clear biological significance when comparing across regions without context of cell or post-synaptic density within the same regions not measured in this study. We chose to include density heatmaps in supplemental figures as to supply additional data for readers with advance knowledge of spinal cord volumetric anatomy. Heatmaps and overlayed dendrogram clustering was analyzed using ComplexHeatmap package (version 2.10.0). Statistical comparisons between cervical levels were compared using a Kruskal-Wallis with subsequent Dunn’s test for multiple comparisons with false discovery rate test correction. P values < 0.05 were considered significant, p values <0.07 are reported on graph. Data were visualized and graphs were generated using custom R scripts, syGlass (syGlass 1.8), and in Prism (Graph Pad Prism 6.0). syGlass volumetric objects were created using ∼200 raw fluorescent MIP images or iLastik pixel probability maps overlayed with atlas regions. Varying Z-voxel sizes were used to create pseudo-MIP across multiple images utilizing the cut tool to isolate registered individual cervical levels.

---

**Supplemental Figure 1: Validation of SpinalTRAQ anatomical precision with classification and quantification of motoneuron somas and dendritic processes in lamina 9.** (A) Representative images of a spinal cord section from a transgenic ChAT-GFP reporter mouse. Left, raw fluorescent images of a whole section (top, scale bar 400um) and enlarged area showing ventral horn region (bottom, scale bar 50um) with GFP+ motoneurons. Corresponding pixel classified images (probability maps) after training to identify motoneuron somas (magenta), dendritic processes (green), white matter (yellow), gray matter (blue), and background (red) are shown at right. (B) Quantification of the percentage of ChAT+ GFP+ somas and (C) dendritic processes found within the Lamina 9 motor pools delineated by the digital atlas. (D) Condensed probability maps showing somatic (magenta) and neuronal process (green) labeling in one representative ChAT-GFP animal for the indicated cervical levels C1 through C8.

**Supplemental Figure 2: Density of the CST synaptic projectome throughout the entire cervical spinal cord of healthy adult mice.** Heatmap of the density of pre-synaptic terminals showing the group mean at each cervical level and lamina, with dendrogram clustering of the lamina based on pre-synaptic terminal density in the contralateral hemicord. Hemicord data are presented on different heat scales as noted.

**Supplemental Figure 3: Density of CST synaptic connections with ChAT+ motoneurons in healthy adult mice.** (A) Mean density of pre-synaptic terminals for each motoneuron pool throughout the cervical cord in both hemicords and clustered using a dendrogram for the motoneuron pools of the contralateral side. (n = 4). (B) eGFP+ pre-synaptic terminals in raw fluorescent images of ventral horn from a healthy adult mouse acrossall eight cervical levels with inset of the corresponding Allen Atlas annotation image. . Presynaptic terminals are shown in green. (C) Maximum intensity projection of a confocal z-stack of a cervical spinal cord section of Sun1-sfGFP animal 4 weeks after AAV1-cre injection. SfGFP positive nuclei are shown in green and all nuclei are stained with DAPI and shown in blue. (D) Individual and merged fluorescent channel images of a spinal cord section from aSun1-sfGFP animal 4 weeks after AAV1-cre injection. SfGFP positive nuclei are shown in green, ChAT immunostaining in red, NeuN immunostaining to label neuronal nuclei is shown in cyan, and DAPI total nuclear staining is shown in blue. Merged multichannel image shown at far right (arrows highlighting sfGFP positive nuclei, scale bar 20um).

**Supplemental Figure 4: Density of reinnervation of the entire denervated hemicord by the contralesional CST synaptic projectome.** (A) Density heatmaps demonstrating the mean of the group at each cervical level and lamina clustered using a dendrogram for the lamina of the sham innervated side. Only the denervated hemicord is shown for the three post stroke groups. (B) Density heatmaps demonstrating the mean of the group at each laminae 9 motor pool of the denervated hemicord annotated in the atlas, clustered using a dendrogram for sham denervated. (Sham: n= 4, 1-Week PS: n = 4, 4-Week PS: n = 5, 6-Week PS: n = 4)

**Supplemental Figure 5: CST synaptic projectome throughout the entire innervated hemicord after stroke.** (A) Heatmap of normalized pre-synaptic terminals demonstrating the mean of the group at each cervical level and lamina clustered using a dendrogram for the lamina of the sham innervated side.

**Supplementary Video 1: Rostral to Caudal Flythrough of a Mouse Cervical Cord Imaged Via STPT Reconstructed in syGlass**

**Supplementary Video 2: Rotational Video of a Mouse Cervical Cord Imaged Via STPT Reconstructed in syGlass**

**Supplementary Video 3: A Flip Through Video of the SpinalTRAQ Cervical Average Template with Atlas Digital Annotations**

**Supplementary Video 4: A Flip Through Video of a Mouse Cervical Cord Registered into SpinalTRAQ Average Template/Volume with Classified White and Gray Matter Supplementary Video 5: A Flip Through Video of a Mouse Cervical Cord Registered into SpinalTRAQ Average Template/Volume with Classified Pre-Synaptic Sites Supplementary Video 6: Rotational Video of a Mouse Cervical Cord ilastik Classified Synapses Reconstructed in syGlass**

## Author Contributions

Conceptualization, K.P., M.K., E.J.P., D.M.O.R, M.P.G; Methodology, K.P, M.K, E.J.P., D.M.O.R, M.P.G., A.N., A.D.A, W.X, X. K.; Software, M.K., A.N., A.D.A, D.M.O.R; Validation, K.P, M.K, A.N., A.D.A, E.J.P., D.M.O.R, W.X, X.K.; Formal Analysis, K.P., M.K., D.B., A.N., A.D.A.; Investigation, K.P., M.K., D.B., A.N., A.D.A.,E.J.P, X.K., D.M.O.R; Writing – Original Draft; K.P., M.K., D.B., D.M.O.R, M.P.G; Writing – Review & Editing, K.P., M.K., D.B., A.N., A.D.A.,W.X.,E.J.P, X.K., D.M.O.R, M.P.G.; Visualization, M.K., A.N.; Supervision, D.M.O.R, M.P.G.; Funding Acquisition, D.M.O.R, M.P.G.

## Declaration of generative AI and AI-assisted technologies

None.

## Acknowledgements

We thank Drs. Helen Lai and Julia Kaiser for their thoughtful manuscript feedback and Drs. Sam Pappas and William Dauer for the gift of ChAT-GFP mice.

