## Supplementary figures and images for "SpinalTRAQ: A novel volumetric cervical spinal cord atlas identifies the corticospinal tract synaptic projectome in healthy and post-stroke mice"

### Supplemental Figure 1

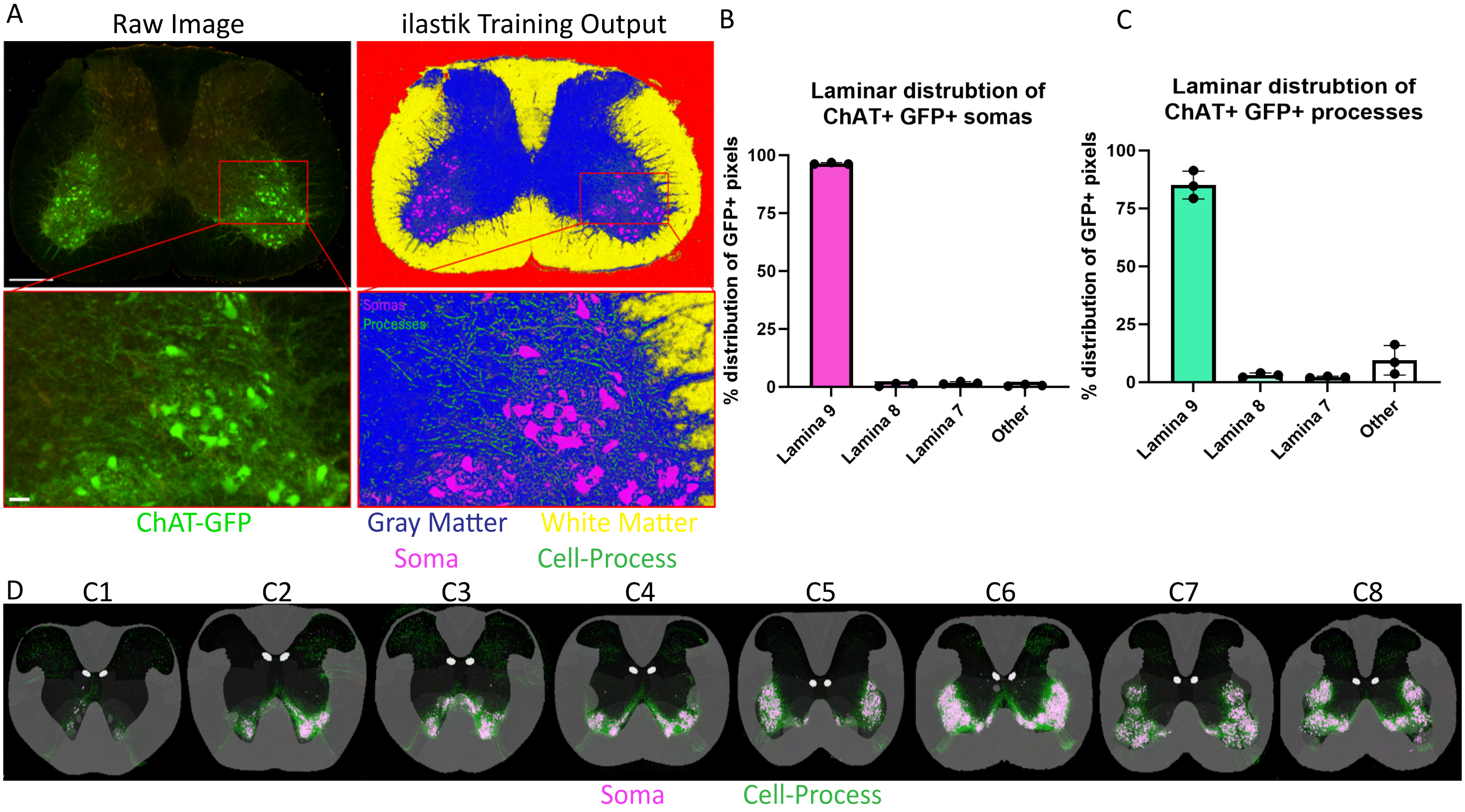

### Supplemental Figure 2

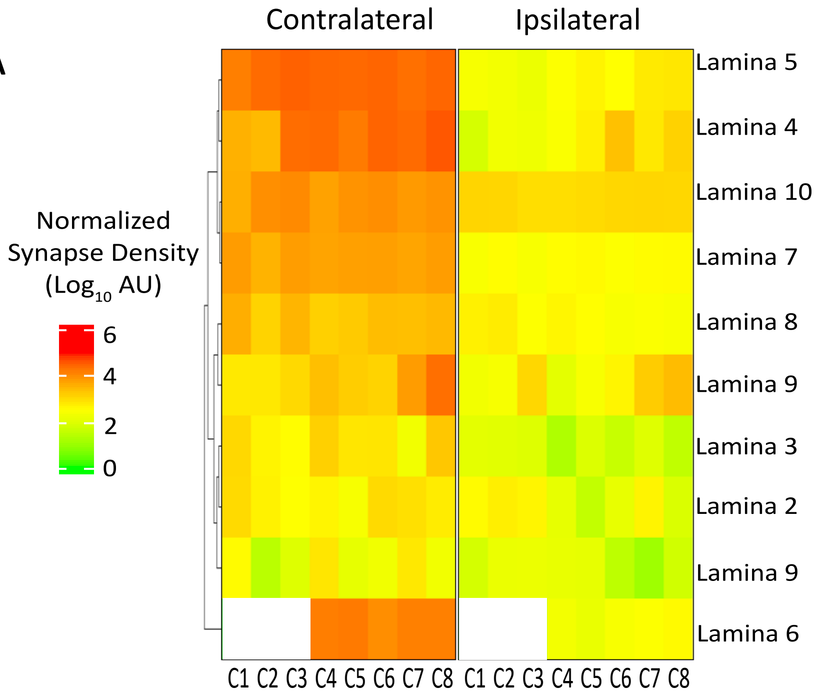

### Supplemental Figure 3

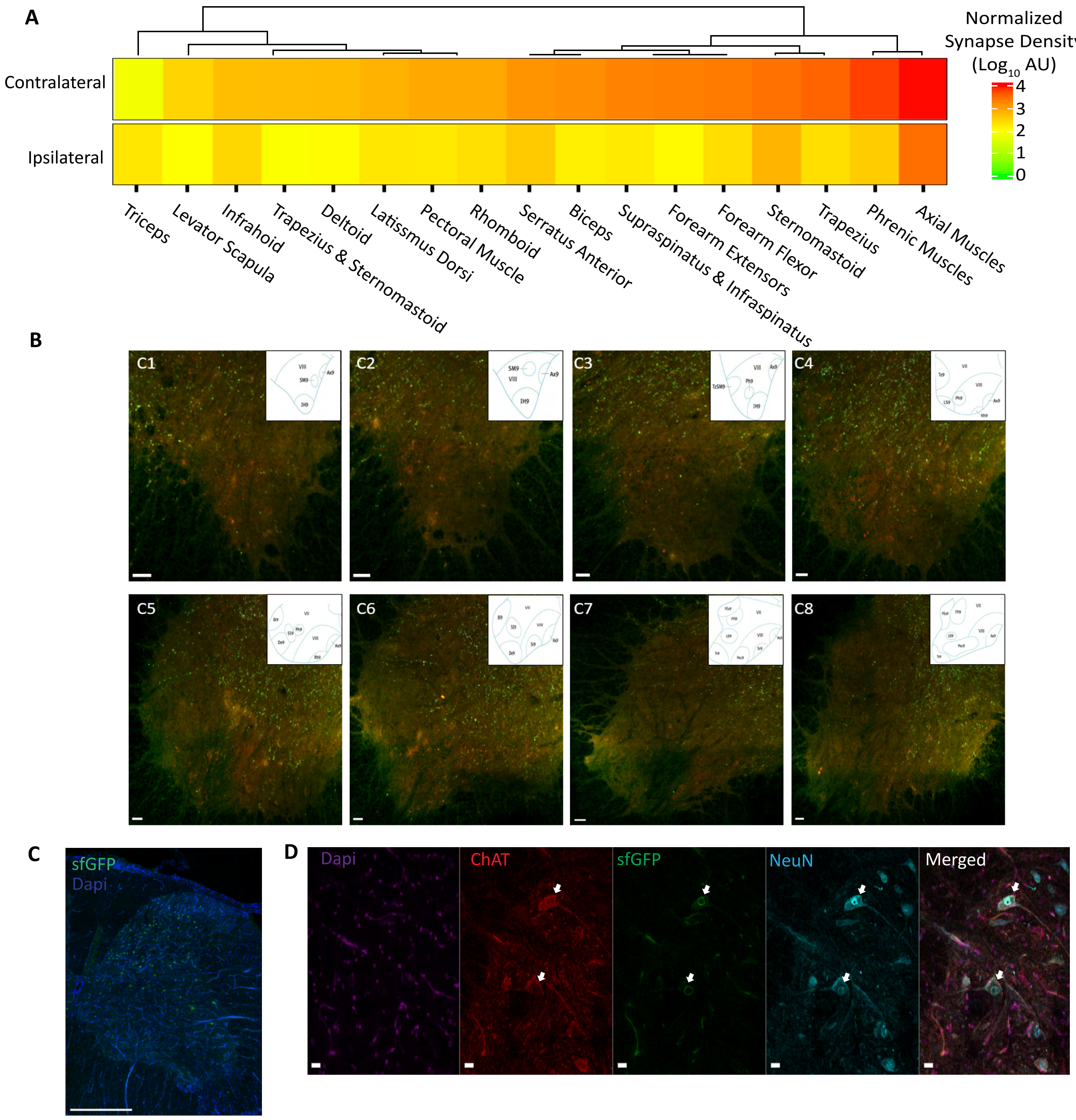

### Supplemental Figure 4

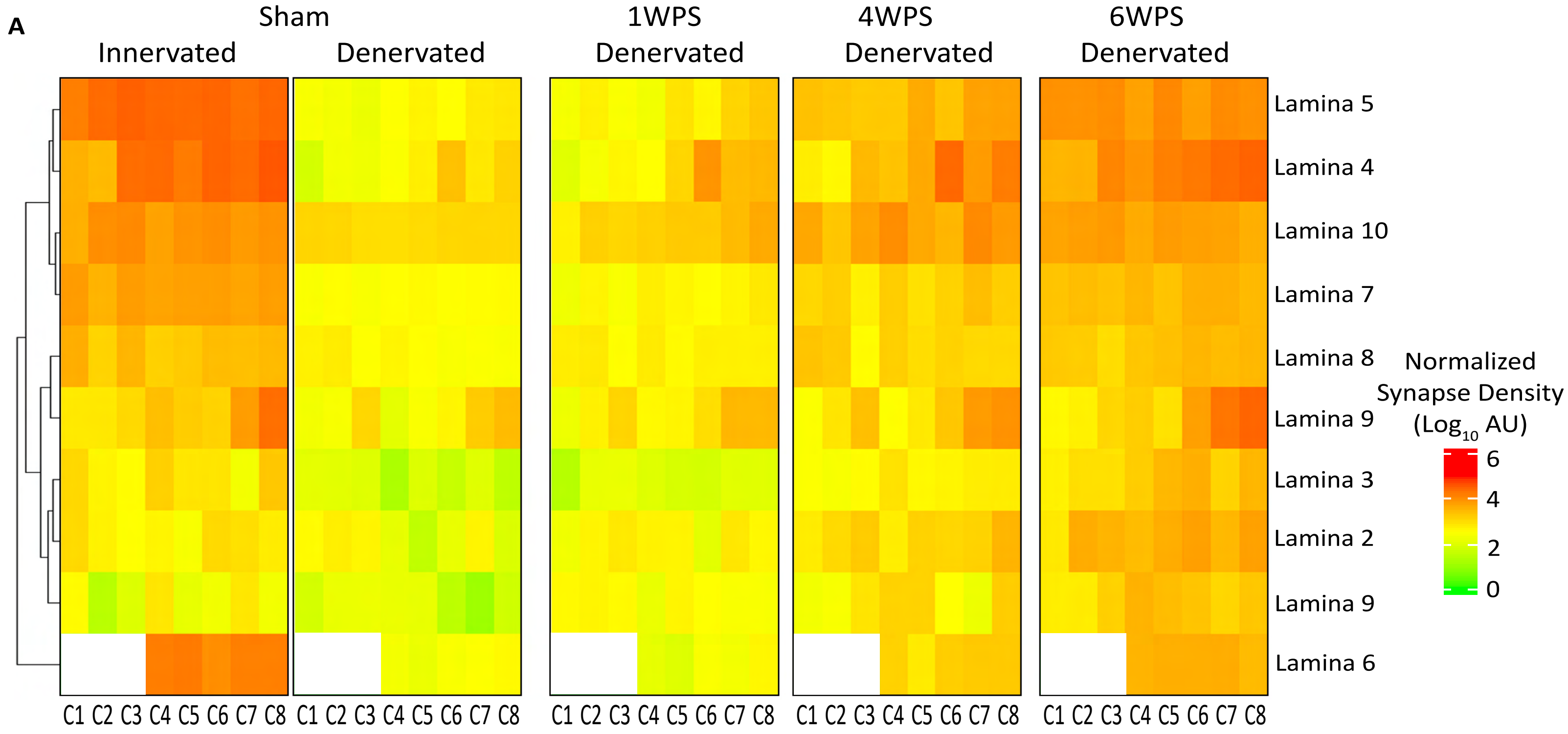

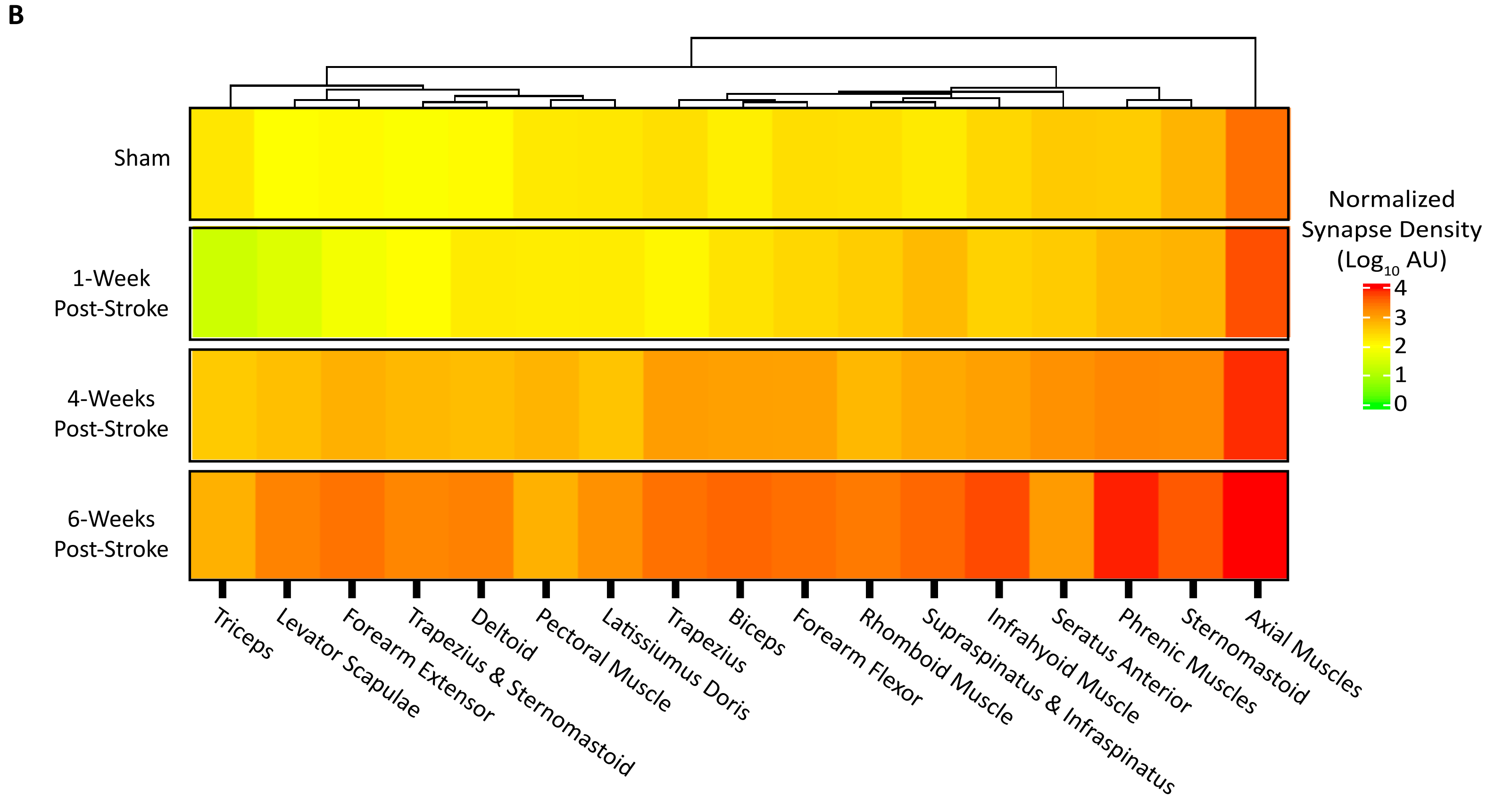

### Supplemental Figure 5

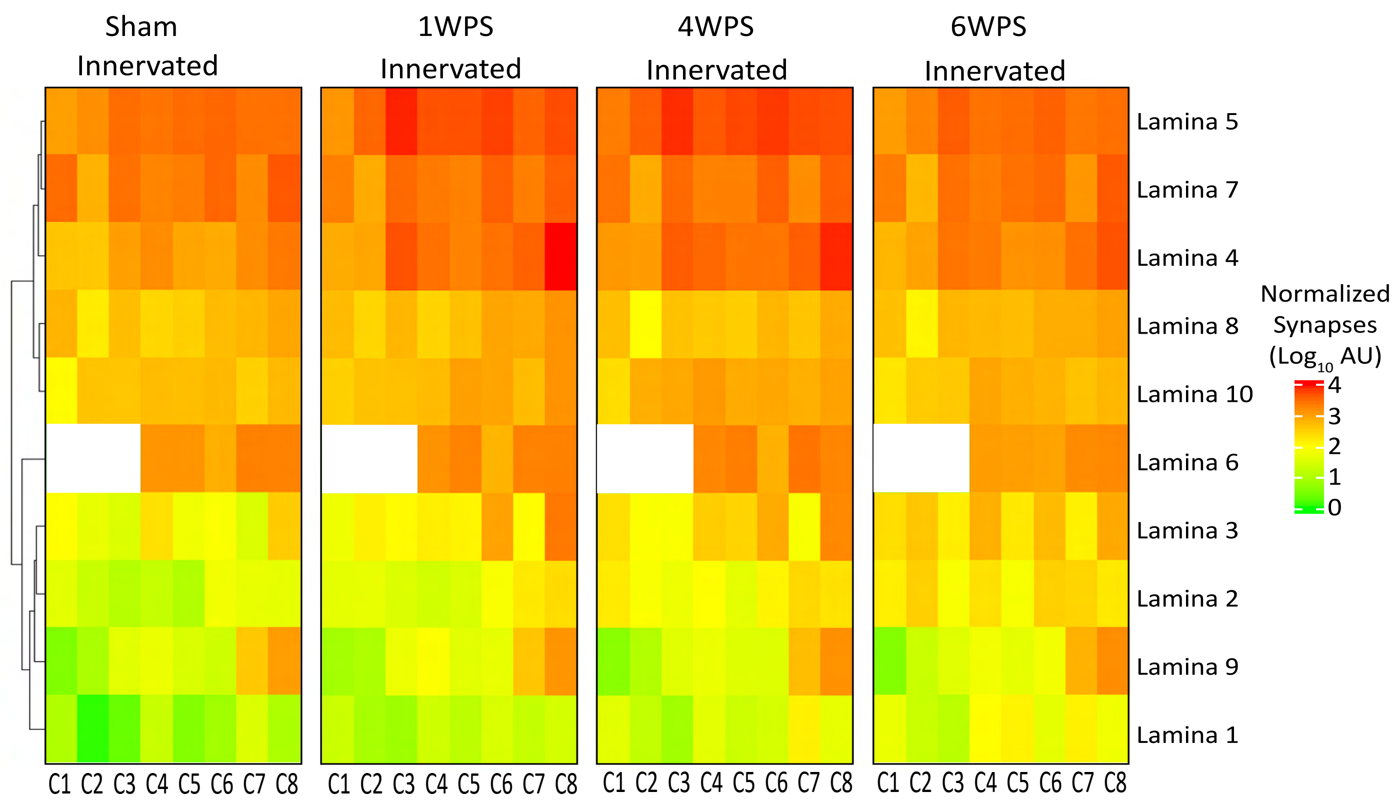
